# C1orf112 is a novel regulator of interstrand crosslink that decreases FIGNL1-RAD51 interaction

**DOI:** 10.1101/2022.10.06.511186

**Authors:** Edgar Pinedo-Carpio, Julien Dessapt, Romain Villot, Lauralicia Sacre, Abba Malina, Jonathan Boulais, Elise G. Lavoie, Vincent Luo, Ana-Maria Lazaratos, Jean-François Côté, Frédérick Mallette, Alba Guarné, Amelie Fradet-Turcotte, Alexandre Orthwein

**Author notes:** co-first authors (Edgar Pinedo-Carpio, Julien Dessapt). co-corresponding authors, Address correspondence to: **Alexandre Orthwein, M.Sc., Ph.D.**, Lady Davis Institute, Segal Cancer Centre, Jewish General Hospital, Montreal, QC, H3T 1E2 Canada, **Amelie Fradet-Turcotte, Ph.D.**, Oncology Division, CHU de Québec Research Center-Université Laval, Cancer Research Center - Université Laval, Quebec, QC, G1R 2J6 Canada.

## Abstract

Interstrand DNA crosslinks (ICLs) represent complex lesions that block essential biological processes, including DNA replication, recombination, and transcription. Several pathways have been involved in ICL repair, in particular nucleotide excision repair (NER), translesion DNA synthesis (TLS), Fanconi anemia (FA), and homologous recombination (HR). Still, the extent of factors involved in the resolution of ICL-induced DNA double-strand breaks (DSBs) remains poorly defined. Using CRISPR-based genome-wide screening, we identified the poorly characterized C1orf112 (also known as Apolo1) as a novel sensitizer to the clinically relevant ICL-inducing agent mafosfamide. Consistently, we noted that low expression of C1orf112 correlates with increased sensitivity to a series of ICL agents and PARP inhibitors in a panel of cell lines. We showed that lack of C1orf112 does not impact the initial recruitment and ubiquitylation of FANCD2 at the ICL site but rather impairs the resolution of RAD51 from ICL-induced DSBs, thereby compromising homology-directed DNA repair pathways. Our proximal mapping of C1orf112 protein neighbours coupled to structure-function analysis revealed that C1orf112, through its WCF motif, forms a complex with the N-terminal domain of the AAA+ ATPase FIGNL1 and regulates the interaction of FIGNL1 with RAD51. Our work establishes the C1orf112-FIGNL1 complex as an integral part of the HR-mediated response to ICLs by regulating the unloading of RAD51 during ICL repair.

## INTRODUCTION

Interstrand crosslinks (ICLs) are highly cytotoxic DNA lesions that produce physical obstacles for vital biological processes, including DNA replication, transcription, and recombination. ICLs can emerge from endogenous reactive aldehydes, but more often occur upon exposure to either naturally occurring compounds (e.g. mitomycin C), or chemically synthesized drugs, such as cisplatin or cyclophosphamide (reviewed in^1^). These ICL-inducing drugs are widely used as chemotherapeutic agents in the treatment of both solid tumours and blood cancers^2^. Aside from compromising essential biological processes, ICLs, if left unrepaired, result in a variety of genomic abnormalities, ranging from point mutations to chromosome breakage or missegregration, up to mitotic catastrophe (reviewed in^3^).

Each type of ICL-inducing agents leads to different types of intra- and interstrand crosslinks that distort the double helix to different levels^1^. To detect, signal and repair these lesions, cells rely on a complex response which typically gets triggered when the replication machinery encounters the ICL, and involves the precise and coordinated effort of multiples pathways, including the Fanconi Anemia (FA), Nucleoride Excision Repair (NER, and the Homologous Recombination (HR) repair pathways, alongside the ATR checkpoint signaling pathways^1^. ICLs are initially recognized by the FANCM complex (FANCM, FAAP24, MHF1 and MHF2) that serves as a platform for the docking of seven additional FANC proteins (FANCA-C, FANCE-G, FANCL) as well as two additional associated proteins (FAAP20 and FAAP100) that altogether form the FA core complex. Together with FANCT (UBE2T), the E3-ubiquitin ligase FA core complex promotes the monoubiquitylation of the effector FANCI/FANCD2 (ID2) heterodimer, a critical step in the processing and subsequent repair of the ICLs. Once assembled and activated at the lesion, the ID2 heterodimer recruits structure-specific nucleases that promote ICL unhooking; and acts as a molecular hub for the recruitment of translesion synthesis (TLS) DNA polymerases which bypass the ICL adduct, and the subsequent recruitment of HR factors involved in the repair of the DNA double-strand breaks (DSBs) generated during this process^1^. The last few years witnessed the discovery of an important decision point during the detection and early processing of ICL, named the ICL repair pathway choice. This decision is made between replication-independent and -dependent pathways and appears to be dictated by the structure and the location of the ICL as well as the phase of the cell cycle in which it is detected ^1^. Recent dissection of the replication-dependent ICL repair pathway has identified a series of novel players such as TRAIP^4,5^, NEIL3^6^, and SCAI-REV3^7,8^ that act before the TLS and HR pathways, respectively.

Aside from a better understanding of ICL repair pathway choice, several reports have highlighted the complexity of the final stages of ICL repair, such as the resolution of RAD51-mediated strand invasion. For instance, the AAA^+^ ATPase FIGNL1 has been shown to bind RAD51 and promote its unloading from DNA^9^, a function that is limited by the SWSAP1-SWS1-SPIDR complex^9,10^. More recently, the helicase HELQ^11^ and the HROB–MCM8–MCM9^12^ complex have been shown to contribute to parallel pathways that promote DNA repair synthesis following D-loop formation. However, the spatiotemporal regulation of these functions is still unknown.

Although the first model of ICL repair was drawn in the early 2000’s, the extent of factors involved in the regulation of this repair pathway remains largely unclear. Here, our CRISPR-based genomic screens identified the poorly characterized C1orf112 as a novel player in the response to ICLs. In-depth characterization delineated its contribution to the regulation of the HR pathway during the ICL response. Proximal mapping of C1orf112 proximity interactors coupled to structure-function validation identified the AAA+ ATPase FIGNL1 as a constitutive partner in the unloading of RAD51 at ICL-induced DSBs. Our model defines the C1orf112-FIGNL1 complex as an integral component of the resolution of ICL-induced DSBs by promoting the dissociation of RAD51.

## RESULTS

### CRISPR screening identifies C1ORF112 as a modulator of ICL repair in BL cells

To better understand the factors influencing ICL repair, we undertook CRISPR/Cas9 dropout screens in two human Burkitt’s lymphoma (BL) cell lines, Namalwa and Raji, using the metabolized version of the ICL agent, cyclophosphamide, as a selective drug (Mafosfamide (MAF)). First, we generated stable Namalwa and Raji cell lines that expressed Cas9 using lentiviral transduction and confirmed genome editing efficiency by targeting the FAM83G gene as previously described^13^. Next, CRISPR-based genome-wide screening was initiated by transducing the TKO v1 sgRNA library in both human BL Cas9-cell lines^14,15^ and transduced cells were amplified under puromycin (2 μg/ml) selection for 7 days (**Fig. 1A**). Both Namalwa and Raji transduced cell lines were subsequently treated with either preoptimized doses of MAF (IC25) (1.30 μM and 1.57 μM, respectively) or with DMSO as a vehicle for 14 doubling time (14 days for both cell lines) before being processed for next-generation sequencing (NGS). The DrugZ algorithm was used to determine the relative abundance of each sgRNA and identify genes whose knockout sensitizes cells to MAF (NormZ score < −2.5, p < 0.005^16^) (**Fig. 1B, S1A and Table 1**). In both screens, pathway enrichment analysis identified ICL repair and HR as two biological pathways statistically enriched among our identified genes of interest (**Fig. S1B, data not shown**). As expected, most factors of the Fanconi Anemia core complex and factors implicated in HR repair emerged as top sensitizers in both cell lines (**Fig. 1B, S1A and Table 1**), validating our CRISPR-based screening approach to identify new genes influencing ICL repair. Of note, the recently characterized *FIGNL1*^9,17^, *MCM8-MCM9*^12^ *and SCAI*^7^ scored significantly on both of our screens (**Fig. 1B, S1A, Table S1)**.

**Figure 1.**
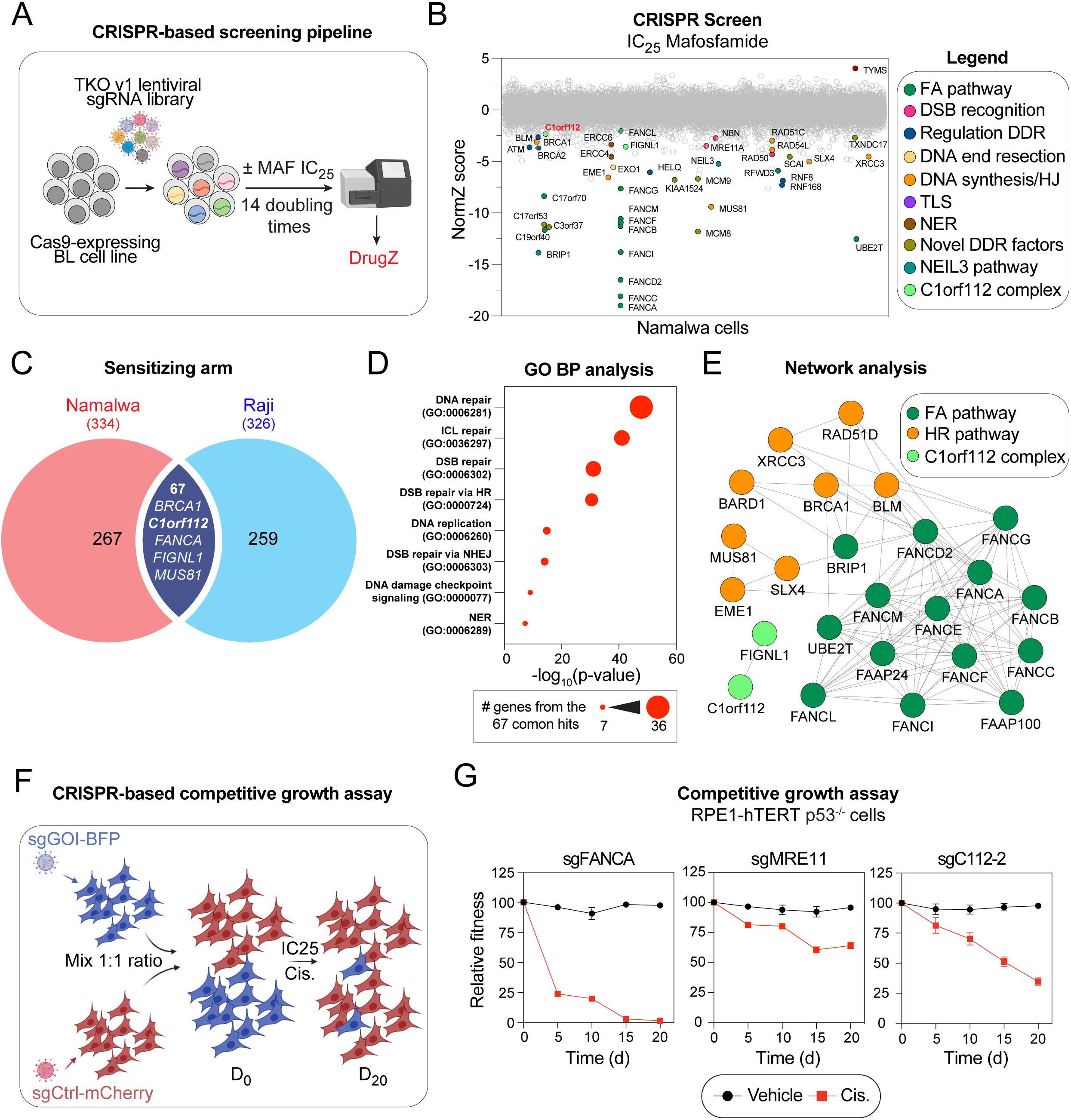
CRISPR-based genome-wide screening identifies C1orf112 as a novel modulator of mafosfamide sensitivity. (A) Schematic representation of the CRISPR-based screening pipeline used in Burkitt lymphoma cells and the screen analysis using the DrugZ algorithm. (B) Horizontal scatter plot of DrugZ-generated ranking of the Namalwa mafosfamide CRISPR screen. NormZ values are plotted on the Y-axis, and gene names are plotted on the X-axis. FA: Fanconi Anemia, DSB: Double-Strand Break, DDR: DNA Damage Response, HJ: Holliday Junction, TLS: Translesion Synthesis, NER: Nucleotide Excision Repair. (C) Venn diagram displaying common sensitizing genes from Namalwa and Raji screens (for genes with NormZ ≤ 2) (D) Gene ontology biological processes (GO BP) diagram obtained from common sensitizing genes from Namalwa and Raji screens. GO term enrichments are ranked by statistical significance (p-value). The size of the circle indicates the number of the 67 common gene hits within a pathway. (E) Network analysis displaying protein physical interactions using Cytoscape and the GeneMANIA package. The C1orf112-FIGNL1 complex is represented in the vicinity of FA and HR repair networks. (F) Schematic representing the pipeline used in the two-color competition assay. RPE1-hTERT p53^-/-^ cells were transduced with a sgRNA targeting either the gene of interest (GOI) coupled with Blue Fluorescent Protein (BFP) or LacZ gene coupled with mCherry. BFP- and mCherry-expressing cells were mixed at a 1:1 ratio at day 0 (D_0_) and treated with IC25 cisplatin (Cis.). BFP-positive over mCherry-positive cells ratio was followed over 20 days (D_20_). (G) Competitive growth results obtained in RPE1-hTERT p53^-/-^ cells targeted with sgRNA against FANCA, MRE11 or C1orf112 (C112). Cells were either treated with DMSO as a vehicle or cisplatin at its IC25 (2.4 μM). Data are represented as the mean ± SEM (n = 3 technical replicates).

We identified 67 target genes that function as sensitizers in both BL cell lines (**Fig. 1C, Table 1**). Gene ontology analysis of these hits revealed enrichment for specific biological processes related to DNA repair, DNA replication and DNA damage checkpoint (**Fig. 1C-D, Table S2**) and the network constructed with these common sensitizers shows high connectivity between genes of the FA and HR pathway (**Fig. 1E and S1C**). According to this network, we found that C1orf112 behaves similarly to factors of the cellular response to ICL agent in BL cell lines. Previous reports identified C1orf112 (aka Apolo1) as a modulator of chromosome segregation through the regulation of PLK1 activity^18^. As no other genes associated with mitosis progression were enriched in our screens, we reasoned that a different function of C1orf112 may be important for the cellular response to ICL.

To validate the C1orf112 contribution in the response to ICL agents, we targeted it using a single-guide RNA (sgRNA) in a twenty-day CRISPR-based growth competition assay^19^ that was performed in hTERT immortalized non-transformed human retinal pigmented epithelial cell line RPE-1 (henceforth termed RPE-hTERT p53^-/-^ cells) (**Fig. 1F, Table S1**). Briefly, stable cell lines expressing a sgRNA of interest along with a BFP-reported or a control sgRNA against LacZ along with a mCherry reporter were established. At D_0_, the cell line expressing the sgGOI-BFP was mixed to a 1:1 ratio with the sgCtr1-mCherry cell line and treated for 20 days (D_20_) with an IC25 of the ICL-inducing agent cisplatin (2.4 nM) before BFP and mCherry measurement at different time points. For comparison, we also tested sgRNAs that target representative genes of the FA (sgFANCA) and the HR pathways (sgMRE11). In this assay, C1orf112 knockout hypersensitizes RPE-hTERT p53^-/-^ cells to cisplatin, demonstrating that the phenotype associated with C1orf112 depletion is neither cell line nor ICL agent specific (**Fig. 1F and S1E**). The effect of C1orf112 depletion on the cellular fitness of cisplatin-treated cells is intermediate of a core FA protein and MRE11 (**Fig. 1F**), suggesting that C1orf112 is important for cellular survival to ICL-inducing agent but may not be a core FA gene. Together, this work demonstrates that our CRISPR-based screening approach is effective for identifying genes that impact the response to ICL agents, including C1orf112 as a new modulator of this response.

### C1orf112 depletion induces genomic instability and senescence in a p53-dependent manner

In a first attempt to define the role of C1orf112 on genomic stability, we depleted it by using small-interfering RNA (siRNA) in Human Bone Osteosarcoma Epithelial Cells (U2OS) cells (**Fig S2A**). At 48 hrs post-transfection, C1orf112-depleted U2OS cells accumulated micronuclei (MNi). MNi formation is triggered by unresolved genomic instabilities, such as DSBs and lagging chromosomes^20^. In C1orf112-depleted cells, we observed that most MNi show negative staining for centromeres and positive staining for the DSB marker g-H2AX (**Fig. 2A-B**), suggesting that they arise from improperly segregated acentric chromosome fragments induced by DSBs^20^. 53BP1 nuclear bodies (NBs) are another marker of genomic instability, representing subnuclear structures assembled around DNA lesions caused by replication stress and transmitted during mitosis to the daughter cells^21^. Interestingly, C1orf112-depleted cells exhibit a 2-fold increase in the number of 53BP1-NBs in the G1 phase of the cell cycle (**Fig. S2B-C**) suggesting that C1orf112 plays a role in preventing replication of stress-induced DNA breaks in mammalian cells. To further explore the impact of C1orf112 on genomic stability, we quantified the number and the intensity of g-H2AX foci that accumulate in interphase of C1orf112-U2OS depleted cells. We found that both the number and the intensity of γ-H2AX were remarkably increased in C1orf112-depleted cells treated with 1 μM of cisplatin for 3 hours compared to the control (cells treated with siCtrl) (**Fig. 2C-D**). Interestingly, depletion of C1orf112 is sufficient to increase the number and the intensity of γ-H2AX in untreated cells, indicative of a contribution of C1orf112 to protect cells from endogenous DNA damage (**Fig. 2C-D**).

**Figure 2.**
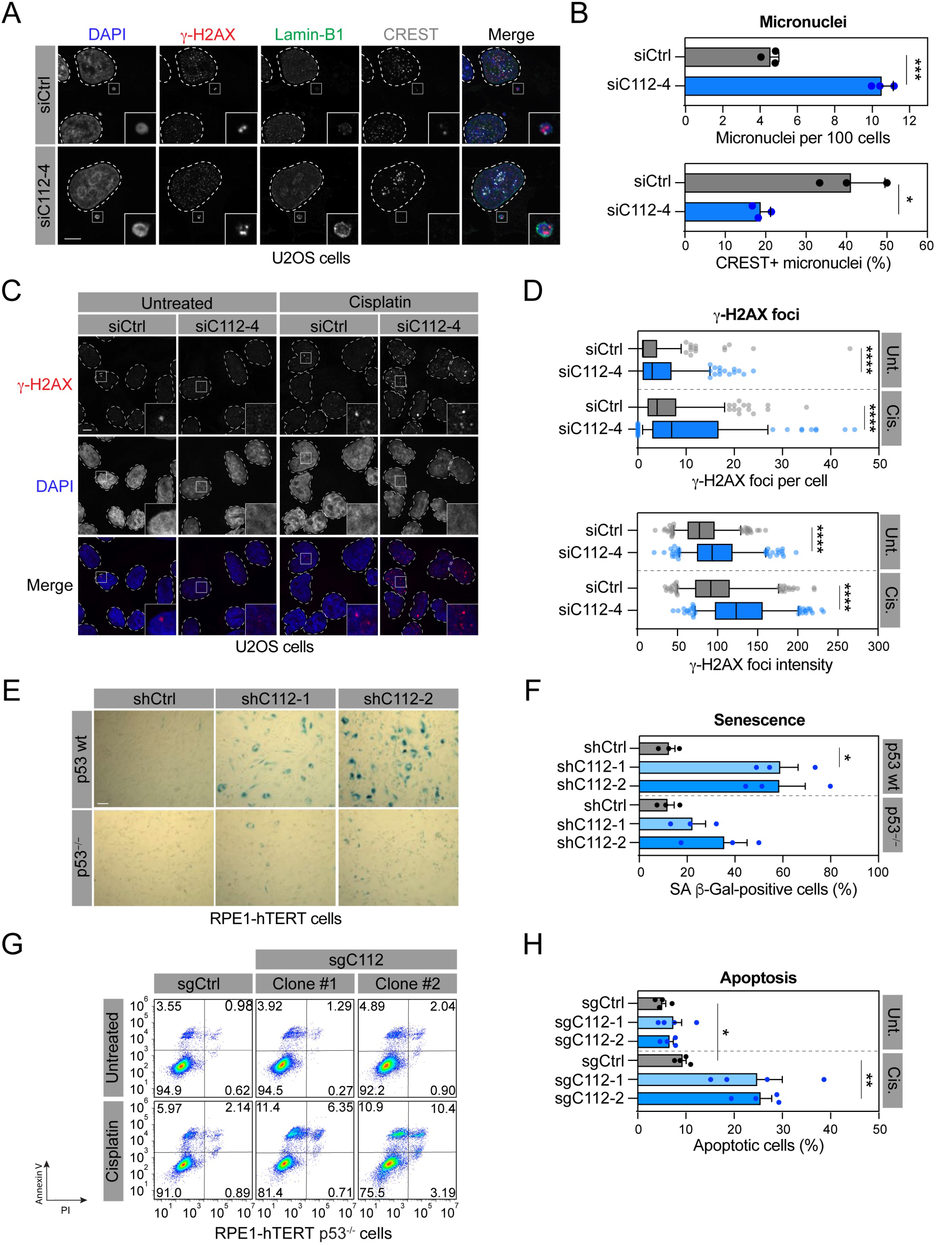
Depletion of C1orf112 lead to genomic instability and senescence-induced cell death. (A) Representative images of U2OS cells depleted or not for C1orf112 and processed for g-H2AX (red), Lamin-B1 (green) and centromeres (CREST) (grey) immunofluorescence. Nuclei were counterstained with DAPI. Scale bar = 5 μm. (B) Quantification of micronuclei Lamin-B1-positive (upper panel) and CREST-positive (lower panel) in cells treated as in (A). Data are represented as the mean ± SD (n = 3 independent experiments). A minimum of 100 cells were analyzed per condition per experiment. Data were analyzed with an unpaired t-test with Welsh’s correction. (C) Representative images of cells depleted or not for C1orf112 and exposed to 1 μM cisplatin or DMSO as a vehicle for 3 hrs. Cells were processed for γ-H2AX (red) immunofluorescence and counterstained with DAPI. Scale bar = 5 μm. (D) Quantification of γ-H2AX foci per cell (top) and γ-H2AX foci intensity (bottom) in cells treated as in (C). Data are represented as the mean ± SD (n = 3 independent experiments). A minimum of 100 cells or foci were analyzed per condition per experiment. Data were analyzed with an unpaired t-test with Welsh’s correction. (E) Representative scatter plot of apoptosis analysis upon treatment with cisplatin in RPE1-hTERT p53^-/-^ cells depleted or not for C1orf112 using shRNA. Apoptosis was induced with pulse treatment of 4 μM cisplatin and compared to vehicle-treated cells (DMSO). Annexin-V/PI staining was conducted 48 hrs post cisplatin treatment and reported as PI content versus Annexin-V intensity in scatter plots. (F). quantification of apoptotic RPE1-hTERT p53^-/-^ cells depleted or not for C1orf112 and treated or not with cisplatin. Data are represented as the mean ± SEM (n = 4, 30,000 cells measured per experiment in each condition). Data were analyzed using a one-way Welch’s ANOVA test and Dunnett’s multiple comparison tests. (G) Representative images of RPE1-hTERT WT and p53^-/-^ cells depleted or not for C1orf112 and stained with senescence-associated (SA) β-Galactosidase. Cells were grown for 24 hrs and stained for SA β-galactosidase. Scale bar = 100 μM (H). Quantification of SA-β-gal positive cells as shown in (G). Data are represented as the mean ± SEM (n = 3 independent experiments). Data were analyzed using a one-way Welch’s ANOVA test and Dunnett’s multiple comparison tests. *p<0.05, **p<0.01, ***p<0.001, ****p<0.0001

In mammalian cells, the presence of DNA damage activates checkpoints that promote cell cycle arrest, senescence or ultimately apoptosis in a p53-dependent manner^22^. Consistent with our previous observations, that C1orf112 depletion results in the accumulation of DNA breaks, we noticed that its depletion by two distinct small-hairpin RNAs (shRNAs) is sufficient to induce senescence in two p53 WT cell lines, RPE1 hTERT and IMR90 (**Fig. 2E-F and S2D-F**). This phenotype was greatly reduced in RPE1 hTERT cells where TP53 was inactivated by CRISPR technology. In fact, all our attempts to generate p53 WT C1orf112 knockout cells line were unsuccessful. In contrast, we were able to isolate two C1orf112^-/-^ clones in RPE1 hTERT p53^-/-^ that we validated by immunoblotting analysis (**Fig. S1G**). In this context, increased levels of apoptosis detected in C1orf112^-/-^ cells by Annexin V/propidium iodide staining showed that both clones are more prone to cell death upon cisplatin treatment (**Fig. 1G, H**), confirming the protective role of C1orf112 against cell death following ICL induction.

### C1orf112 acts downstream of the FA pathway during ICL repair

The response to ICLs relies on the FA pathway to detect covalently linked DNA strands and promote the recruitment of specific nucleases that release the ICL from one of the two parental strands^23^. DSBs left behind by nucleolytic processing are then repaired by homology-directed recombination pathways such as HR and SSA (**Fig. 3A**). To delineate at which step(s) C1orf112 is participating in the ICL response, we first assessed its involvement in the recruitment and the activation of FANCD2 at ICLs. As expected, both FANCD2 accumulation and ubiquitylation at ICLs were abrogated upon depletion of FANCA in U2OS cells, serving as a control (**Fig. 3B-D**). In contrast, C1orf112 depletion had a limited impact on the detection of FANCD2 at g-H2AX foci (**Fig. 3B-C**), and on its mono-ubiquitylation post-cisplatin treatment (**Fig. 3D**). These data suggest that C1orf112 plays a role downstream of the FA pathway during the response to ICLs. In fact, we noticed that FANCA-depleted cells were hypersensitive to both ICL agents MAF and cisplatin in our viability assay, unlike C1orf112-targeted cells, which displayed a much milder response to these drugs (**Fig S3G**).

**Figure 3.**
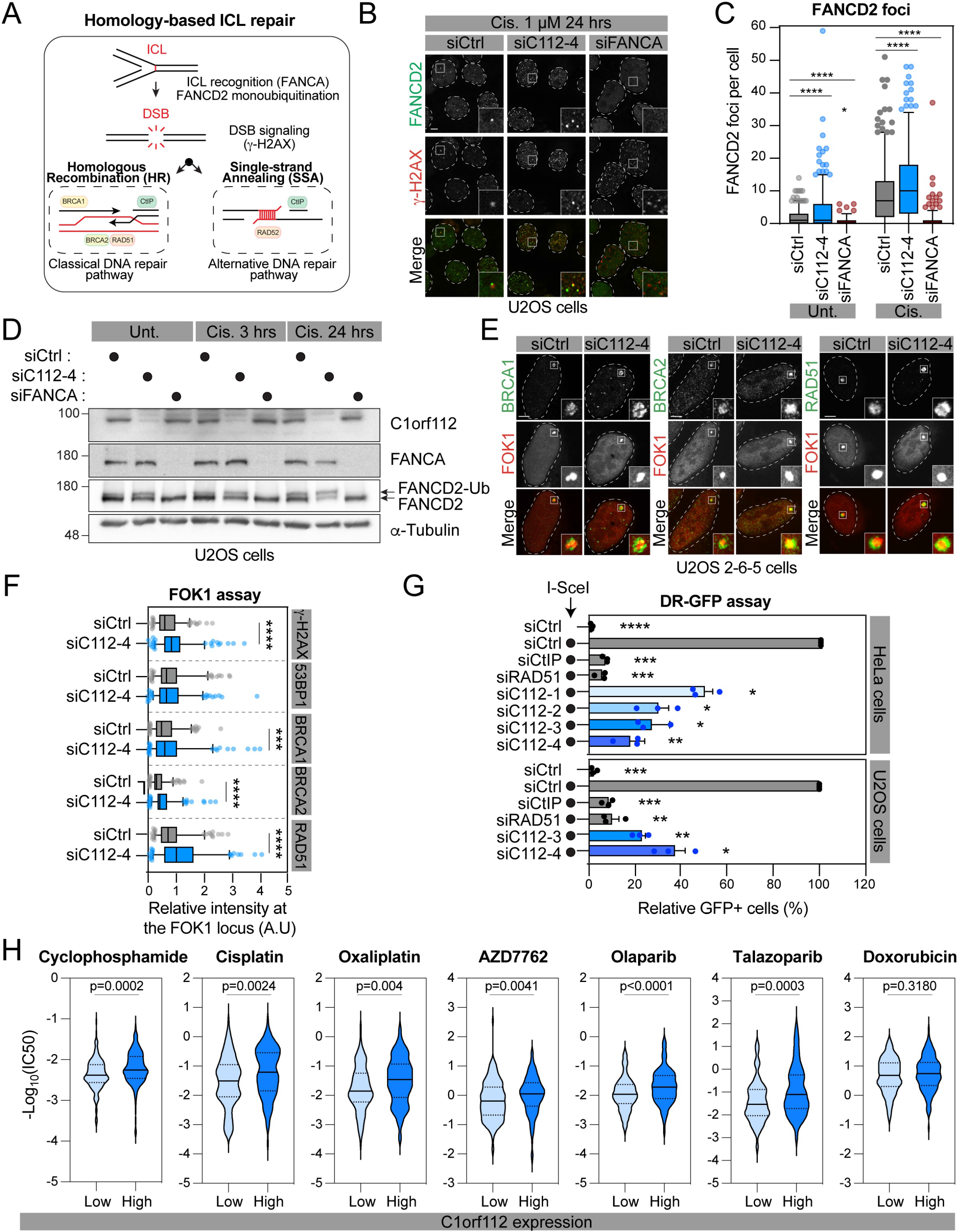
C1orf112 is not a core FA gene but is required for HR. (A) Schematic of ICL repair pathways in mammalian cells. ICL: Interstrand Crosslink. (B) Representative images of U2OS cells depleted or not for C1orf112 or FANCA. Cells were treated with the indicated siRNA for 48 hrs and treated or not with 1 μM cisplatin for 24 hrs before fixation. Cells were processed for g-H2AX and FANCD2 immunofluorescence. Scale bar = 5 μm. (C) Quantification of FANCD2 foci per cell in cells depleted or not for C1orf112 or FANCA. Data are represented as the mean ± SD (n = 3, 100 cells quantified per experiment in each condition). Data were analyzed using a one-way Welch’s ANOVA test and Games-Howell’s multiple comparison tests. (D) U2OS cells were depleted or not for C1orf112 or FANCA and treated with cisplatin or DMSO. Whole Cell Extracts (WCE) were analyzed by immunoblot with anti-C1orf112, anti-FANCA or anti-FANCD2 antibodies. Anti-β-tubulin was used as a loading control. FD2: FANCD1, FD2-ubi: FANCD2 ubiquitinated. (E) U2OS 2-6-5 cells were depleted or not for C1orf112 (for 48 hrs and ER-mCherry-LacR-FOK1-DD expression was induced for 4 hrs before fixation. Cells were then processed for BRCA1, BRCA2 or RAD51 immunofluorescence. Scale bar = 5 μm. (F) Quantification of the ratio of indicated protein foci intensity over ER-mCherry-LacR-FOK1-DD foci intensity as shown in (E). Data are represented as the mean ± SD (n = 4 independent experiments, 50 foci quantified per experiment in each condition). Data were analyzed with an unpaired t-test with Welsh’s correction. (G) Quantification of GFP-positive HeLa and U2OS DR-GFP cells depleted or not CtIP, RAD51 or C1orf112. Twenty-four hours after transfection of the indicated siRNA, cells were transfected with the I-SceI expression plasmid (●) or an empty vector. Data are represented as the mean ± SEM (n = 3 independent experiments, 30,000 cells quantified per experiment in each condition). Data were analyzed using a one-way Welch’s ANOVA test and Dunnett’s multiple comparison tests. (H) Drug sensitivity analysis using expression data for C1orf112 from the CCLE repository-DepMap and IC50 values using the GDSC database. Drug sensitivity of high (dark blue) and low (light blue) C1orf112 expressing cells is shown. Data were analyzed with an unpaired t-test (Cisplatin, Oxaliplatin, Olaparib & Doxorubicin) or a Mann-Whitney test (Cyclophosphamide, AZD7762 & Talazoparib) based on data normality assessed with a Shapiro-Wilk test. *p<0.05, **p<0.01, ***p<0.001, ****p<0.0001

As for g-H2AX foci (**Fig. 2C-D**), we noted that more FANCD2 foci accumulate in C1orf112 depleted cells (**Fig. 3B-C**). To further dissect the stage at which C1orf112 influences the ICL response, we focused our attention on key HR factors and their accumulation at DSBs in absence of C1orf112. We took advantage of a DSB reporter system in which the recruitment of these factors can be quantified at a single genomic locus. Briefly, DSBs are rapidly induced by the recruitment of the ER-mCherry-LacRnls-FOK1-DD fusion protein at a *LacO* array integrated on chromosome 1p3.6 in U2OS cells (U2OS 2-6-5 cell line)^24,25^. In this assay, depletion of C1orf112 correlated with an exacerbated accumulation of several HR factors at FOK1-induced DSBs, including BRCA1, BRCA2 and RAD51 (**Fig. 3E-F and S3A-B**). Of note, C1orf112 depletion did not affect the recruitment of the NHEJ factor 53BP1, suggesting that C1orf112 is primarily involved in HR-mediated processes. To directly assess this question, we used well-established HeLa and U2OS DR-GFP reporter cell lines^26^, as well as the U2OS SA-GFP reporter cell line^27^ to measure DNA repair by HR and single-strand annealing (SSA), respectively (**Fig 3G and Fig. S3C-E**). Depletion of C1orf112 with a series of siRNAs resulted in a marked decrease in both assays, particularly with siRNA-3 and −4, which results in efficient depletion of the protein as determined by immunoblotting analysis (**Fig S3C**). Of note, C1orf112 depletion had a much milder impact in both the DR- and SSA-GFP assays, in comparison to CtIP or RAD51 depletion, suggestive of a regulatory rather than a core contribution of C1orf112 in homology-directed DNA repair pathways.

Consistent with these observations, expression of C1orf112 correlated with the cellular response to ICL-inducing agents in the Genomics of Drug Sensitivity in Cancer Project dataset (CCLE repository-DepMap), where cells expressing higher levels of C1orf112 are more resistant to ICL-inducing agents (**Fig 3H**). Interestingly, high levels of C1orf112 also protect cells from other genotoxic agents that indirectly lead to the formation of DSBs: the Poly [ADP-ribose] polymerase 1 inhibitor (PARPi) Talazoparib and ATR inhibitor (AZD7762). In contrast, higher expression of C1orf112 is not an advantage in treatment that directly induces DSBs such as the topoisomerase II poisoning agent doxorubicin (**Fig 3H**). Chemogenomic profiling of CRISPR screens performed in RPE1 hTERT p53^-/-^ cells with different genotoxic agents showed that C1orff112 depletion impairs predominantly cell survival (NormZ < −2.3) upon exposure to agents that induce replication roadblocks (e.g. cisplatin), base alkylation (MNNG), and oxidative damage (KBrO3, H_2_O_2_) (**Fig S3F, Table S3**)^28^. Altogether, these results support a model where C1orf112 is specifically involved in the repair of DSBs indirectly generated by a subtype replication stress induced by replication fork roadblocks.

### C1orf112 interacts and acts with FIGNL1 in ICL repair

To gain insight into the role of C1orf112 in DNA repair, we mapped its proximal interactome by taking advantage of the miniTurboID technology (Branon et al, 2018a). Briefly, C1orf112 was N-terminally tagged with the promiscuous biotin ligase miniTurbo (mTurboID) and stably expressed in HEK293 Flp-In™ cells (**Fig S4A**). This construct was validated for its ability to efficiently biotinylate proximal partners by immunoblotting analysis (**Fig S4B**). The proximity interacting network of C1orf112 was generated in both untreated cells and cells treated with the radiomimetic drug Neocarzinostatin (NCS) for 3 hrs. A total of 153 high-confidence C1orf112 interactors found in 3 technical replicates were identified (**Table S4**). To pinpoint the interactors that are functionally relevant to the role of C1orf112 in DNA repair, we intersected the list of high-confidence prey obtained in mTurboID (BFDR < 0.01, Saint score = 1) with the NormZ scores obtained from Namalwa and Raji screen datasets (**Fig 4A**). Using this approach, we identified 4 interactors, 3 of which were identified in both treated and untreated cells suggesting that they are constitutive partners of C1orf112 (**Fig 3B**). Two of these interactors, the Pericentriolar material 1 protein (PCM1) and the DNA excision repair protein ERCC6-like (ERCC6L, also known as PICHs), play a role in mitosis where they respectively contribute to the proper assembly of functional centrosomes and the resolution of anaphase bridges^30,31,33^. Therefore, we reasoned that they most likely play a role in the mitotic function of C1orf112 (APOLO1)(Xu et al, 2021). Interestingly, the AAA+ ATPase Fidgetin Like 1 (FIGNL1) was also identified as a constitutive interactor of C1orf112 in our mTurboID. Although FIGNL1 has also been detected at centromeres, two studies support a role for the protein in regulating RAD51 filament during HR repair^9,17^ The interaction between FIGNL1 and C1orf112 has been previously detected with two orthogonal approaches (Yeast two-hybrid^32^, AP-MS^17,34^) which strengthen our findings.

**Figure 4.**
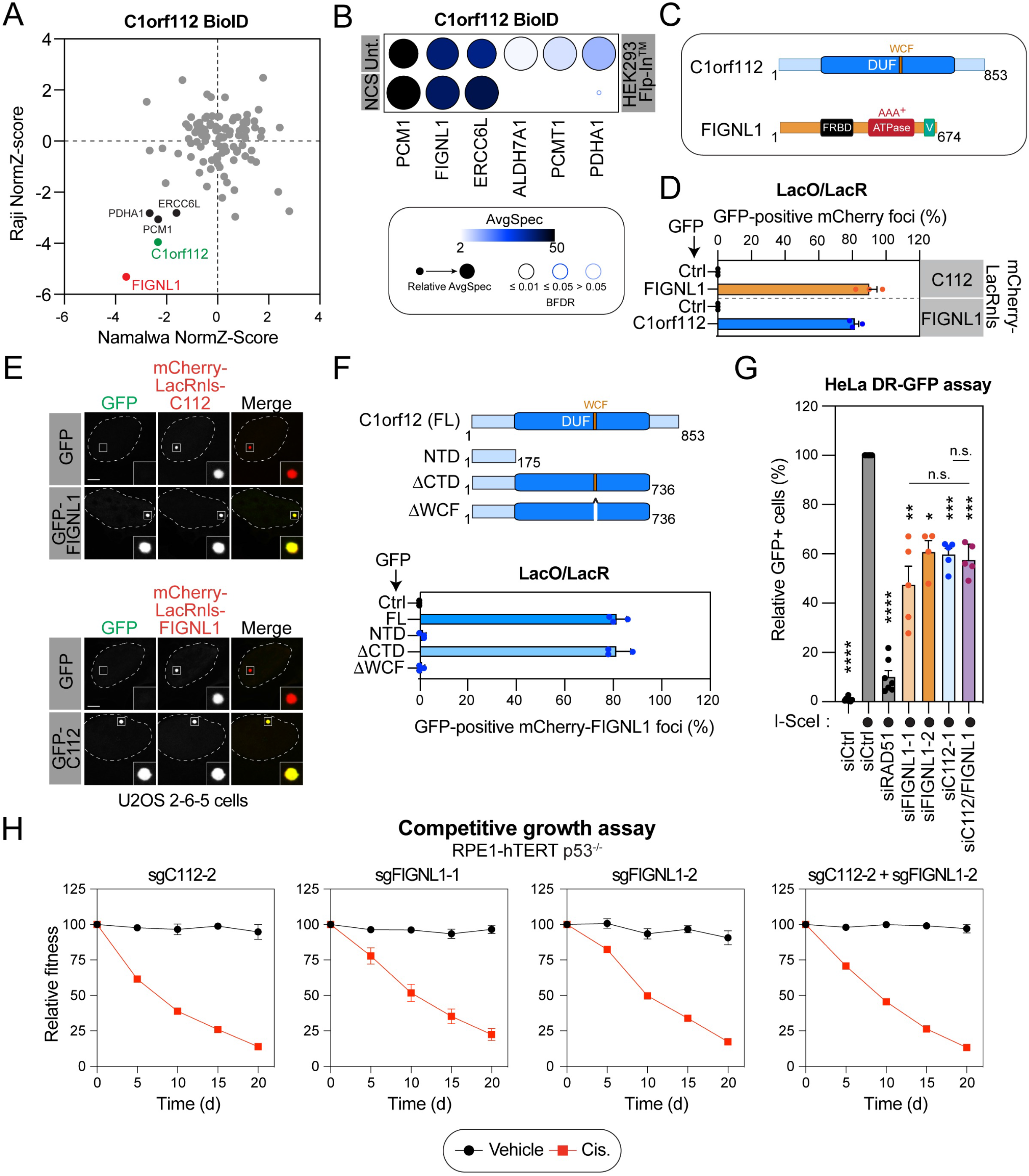
FIGNL1 is a functional interactor of C1orf112. (A) Functional validation of mturboID data using Namalwa and Raji screen datasets. Scatter plots of preys obtained in mturboID proteomic (Fold-change ≤ 1.5) intersect with normalized Z (NormZ) values obtained from both CRISPR screens. High-confidence preys (BFDR < 0.01 & Saint score = 1) behaving like C1orf112 are highlighted. (B) High-confidence proximal interactors of C1orf112 identified by mturboID with or without DNA damages induced with neocarzinostatin (NCS). The spectral counts for each indicated prey protein are shown as AvgSpec. The circle size represents the relative abundance of preys over baits, and node edge color corresponds to the Bayesian False Discovery Rate (BFDR). (C) Schematic representation of C1orf112 and FIGNL1 protein domains. DUF: Domain of Unknown Function. WCF: WCF tripeptide sequence. FRBD: FIGNL1’s RAD51 Binding Domain. V: vps4 domain. (D) Quantification of the colocalization of C1orf112 and FIGNL1 at a *LacO* array. U2OS 2-6-5 cells were transfected with the indicated mCherry-LacRnls and GFP constructs. GFP alone was used as a negative control (Ctrl). Twenty-four hours post-transfections, cells were fixed and mounted for confocal analyses. Data are represented as the mean ± SD (n = 3 independent experiments, 50 cells quantified per experiment in each condition). (E) Representative images of data quantified in (D). Scale bar = 5 μm. (F) Schematic representation of C1orf112 full length (FL) and truncated proteins (upper panel). NTD: N-Terminal Domain. CTD: C-Terminal Domain. Quantification of GFP-tagged proteins colocalizing with mCherry-LacRnls-FIGNL1 in U2OS 2-6-5 cells is presented (lower panel). Cells were transfected and processed as in (D). Data are represented as the mean ± SD (n = 3 independent experiments, 50 cells quantified per experiment in each condition). Data were analyzed with an unpaired t-test with Welsh’s correction. (G) Quantification of GFP-positive HeLa DR-GFP cells depleted or not for RAD51, FIGNL1 and/or C1orf112. HeLa cells containing a DR-GFP reporter cassette were transfected and analyzed as described in Figure 3G. (●) represent I-SceI transfected cells. Data are represented as the mean ± SEM (n = at least 5 independent experiments, 30,000 cells quantified per experiment in each condition). Data were analyzed using a one-way Welch’s ANOVA test and Dunnett’s multiple comparison tests. (H) Competitive growth results in FIGNL1 and/or C1orf112 depleted RPE1-hTERT p53^-/-^ cells. Cells were assayed and treated as in Figure 1F and G. For dual targeting analysis, RPE1-hTERT cells were transduced with a colorless-sgRNA-non targeting control (LacZ), a sgC1orf112 coupled with BFP- and a sgFIGNL1 coupled with mCherry. Data are represented as the mean ± SEM (n = 3 technical replicates). *p<0.05, **p<0.01, ***p<0.001, ****p<0.0001

To validate and further characterize the interaction between FIGNL1 and C1orf112, we took advantage of the U2OS 2-6-5 cell line that contains a *lacO* array integrated into a single locus. In this context, transiently transfected mCherry-LacRnls-bait fusion proteins accumulate at the array in U2OS 2-6-5 cells, and the colocalization of preys is quantified at the mCherry locus. Of note, no FOK1 nuclease is expressed in these experiments. When fused to mCherry-LacRnls, both C1orf112 and FIGNL1 were able to respectively trigger the accumulation of GFP-tagged FIGNL1 and C1orf112 at the LacO array (**Fig 4C-E and S4C-D**). In contrast, neither of the mCherry-LacRnls-tagged proteins recruited GFP alone, which was used as a negative control, at the LacO array. C1orf112 is a protein of 853 amino acids that contains a domain of unknown function (DUF) and a highly conserved WCF tripeptide motif (**Fig 4C**)^32^. Structure-function analysis of C1orf112 truncation and deletion mutants in this experiment revealed that the conserved WCF tripeptide motif is essential for the interaction while the C-terminal region of C1orf112 is not required (**Fig 4F and S4E-F**). Next, we used the DR-GFP reporter assay to investigate the functional relevance of the interaction in DNA repair. Both C1orf112 and FIGNL1 were depleted using siRNA, and the efficiency of the siRNA for FIGNL1 was verified by RT-PCR (**Fig S4G**). As reported in Fig 3, depletion of C1orf112 led to a mild but significant defect in HR repair when compared to the positive control siRAD51 (40 % vs 90 % decrease) (**Fig 4G**). Interestingly, both siRNAs targeting FIGNL1, as well as a double knockdown of FIGNL1 and C1orf112, lead to a similar defect in HR than C1orf112 with 50 %, 40 % and 40% decrease, respectively. This result supports a model where functions of FIGNL1 and C1orf112 are functionally epistatic in the HR pathway. In support of this hypothesis, FIGNL1-depleted and FIGNL1-C1orf112-depleted cells exhibit similar sensitivity to cisplatin in CRISPR-based growth competition assay (**Fig 4H**). Furthermore, the depletion of either protein in U2OS cells specifically reduced cell viability in the presence of ICL-inducing agents such as cisplatin, formaldehyde, mitomycin C, chronic exposure to hydroxyurea and PARPi (**Fig S4H**). Altogether, these findings suggest that C1orf112 and FIGNL1 act as a complex in the resolution of DNA breaks that are indirectly created by replication stress.

### C1orf112 regulates FIGNL1 interaction with RAD51 and RAD51 resolution at ICLs

FIGNL1 is a protein of 674 amino acids with three conserved domains, i) a RAD51 binding domain (FIGNL1’s RAD51 binding domain, FRBD), ii) an AAA^+^ ATPase domain, and iii) a C-terminal Vps4 domain^9,17^. Interestingly, FIGNL1 has been shown to promote RAD51 dissociation from ssDNA^9,17^. To gain insight into how C1orf112 and FIGNL1 may collaborate during ICL repair, we first took advantage of our ability to detect the interaction of GFP-C1orf112 with mCherry-LacRnls-FIGNL1 in the LacO/LacR assay (**Fig 4D**) to rapidly screen a panel of FIGNL1 truncation and mutant proteins (**Fig 5A**). mCherry-LacRnls-FIGNL1 constructs were designed to remove or abolish the function of the previously described FIGNL1’s RAD51 binding domain (FRBD)^17^ and AAA^+^ ATPase activity of FIGNL1^9^ (**Fig 5A and S5A**). In this experiment, FIGNL1 constructs lacking the FRBD (ΔFRBD) or the ability to interact with RAD51 (F295E) still efficiently recruited GFP-C1orf112 to the LacO array (**Fig 5B, Fig S5B-C**). Similarly, none of the single (K447A and D500A) and double (KDm: K447A/D500A) mutations of residues that are highly conserved within the AAA^+^ ATPase domain impacted the interaction of FIGNL1 with C1orf112 (**Fig 5B, Fig S5B-C**). Consistent with the model that RAD51 is not required for the interaction, similar levels of GFP-C1orf112 were detected at the mCherry-LacRnls-FIGNL1 in cells treated with siRNA control or against RAD51 (**Fig S5D-E**). Interestingly, the fact that FIGNL1 constructs encompassing amino acids 1-120 or 121-674 are unable to recruit C1orf112 to the array, suggest that the C1orf112 binding motif (C112BM) of FIGNL1 spans over the intersection of these constructs (**Fig 5A**). Consistently, the FIGNL1 fragment 1-360 efficiently promoted the accumulation of C1orf112 in our assay.

**Figure 5.**
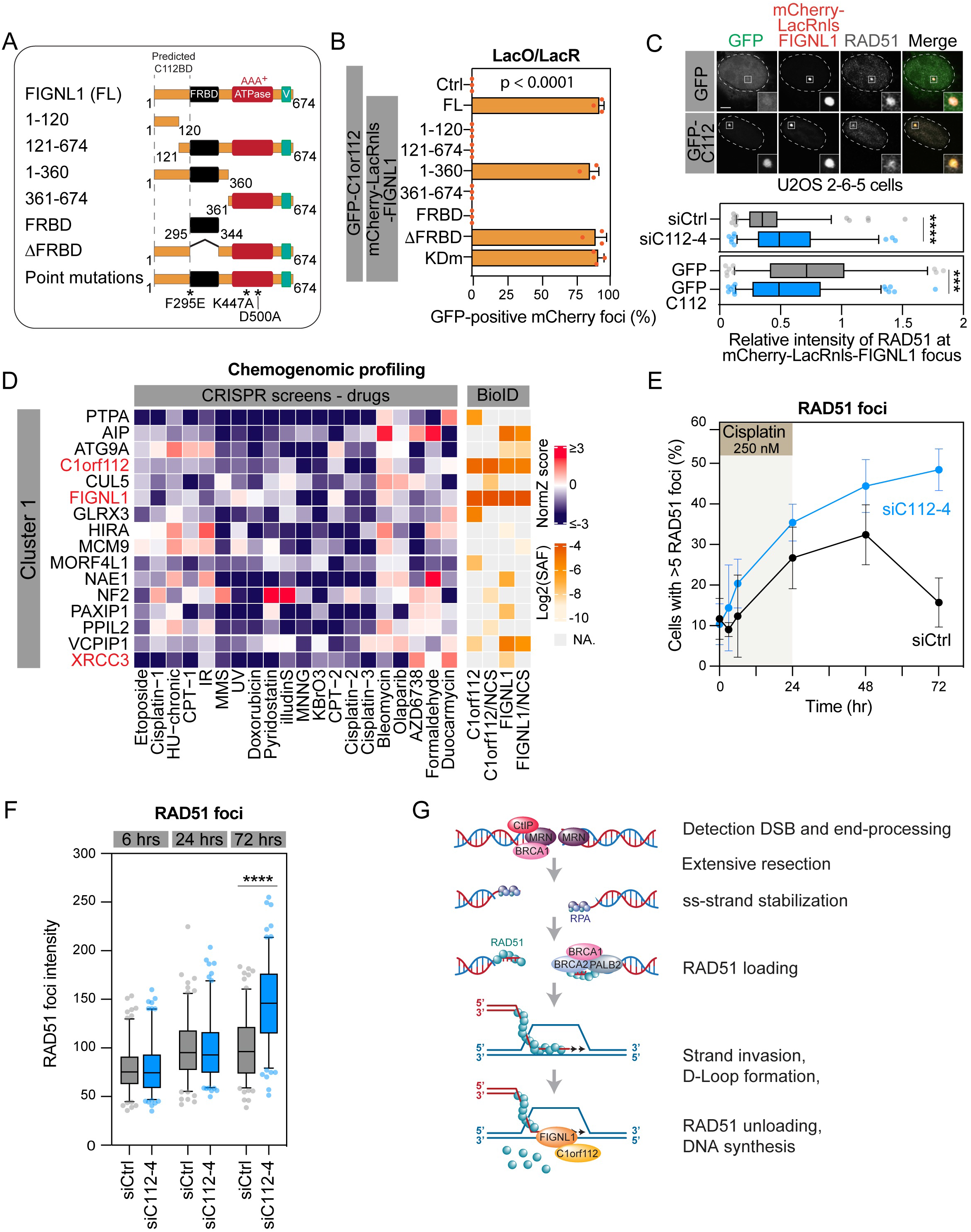
The C1orf112-FIGNL1 complex regulates RAD51 unloading in HR. (A) Schematic representation of FIGNL1 full length (FL), proteins fragments and mutant proteins used in this study. (B) Quantification of mCherry-LacRnls constructs colocalizing with GFP-C1orf112 in U2OS 26-5 cells. Cells were treated and processed as described in Figure 4D. Data are represented as the mean ± SD (n = 3 independent experiments, 50 cells quantified per experiment in each condition). Data were analyzed with an unpaired t-test with Welsh’s correction. (C) Upper panel: Representative images of U2OS 2-6-5 cells transfected with mCherry-LacR-FIGNL1 and GFP-C1orf112. GFP was used as a negative control. Fixed cells were processed for RAD51 (grey) immunofluorescence. Scale bar = 5 μm. Lower panel: Ratio of the intensity of RAD51 over mCherry signal at the *LacO* array foci. Cells were either transfected with mCherry-LacR-FIGNL1 alone or with a GFP-tagged construct 24h before fixation. When indicated, cells were treated with siRNA 24 hrs prior to plasmid transfection. Data are represented as the mean ± SD (n = 3 independent experiments, 50 foci quantified per experiment in each condition). Data were analyzed with an unpaired t-test with Welsh’s correction. (D) Left panel: Chemogenomic profiling using the NormZ-score from the CRISPR screens performed in RPE1 hTERT Cas9 p53^-/-^ cells with the indicated DNA damaging agents as detailed in^28^. Right panel: Heatmap indicating proximity interaction network of C1orff112 and FIGNL1, treated or not with NCS, with the protein expressed from the indicated gene (Log2SAF: spectral abundance factor). (E) Quantification of U2OS cells with ≥ 5 RAD51 foci following treatment with cisplatin. Cells were depleted or not for C1orf112 prior to treatment with 250 nM cisplatin (t = 0). Twenty-four hours later, cells were washed with PBS and left for recovery. Cells were fixed and then processed for RAD51 immunofluorescence for every time point. Data are represented as the mean ± SD (n = 3 independent experiments, 50 cells quantified per experiment in each condition). Data were analyzed with an unpaired t-test with Welsh’s correction. (F) Quantification of the intensity of RAD51 foci analyzed in (E). Data are represented as the mean ± SD (n = 3 independent experiments, 50 foci quantified per experiment in each condition). Data were analyzed with an unpaired t-test with Welsh’s correction. (G) Proposed model of the FIGNL1-dependent role of C1orf112 in RAD51 unloading following DNA strand invasion. ss: single strand. ***p<0.001, ****p<0.0001

In the LacO/LacR assay, mCherry-LacRnls-FIGNL1 efficiently recruits RAD51 to the array (**Fig 5C, Fig S5F**). Interestingly, we observed a significant increase of RAD51 intensity at the mCherry locus in C1orf112 depleted U2OS 2-6-5 cells (**Fig 5C**), suggesting that C1orf112 negatively regulates the interaction of FIGNL1 with RAD51. Consistent with this hypothesis, ectopic expression of GFP-C1orf112 had the opposite effect and resulted in a reduced signal of RAD51 at the LacO array when compared to the expression of GFP alone (**Fig 5C**). These findings support a model where the C1orf112-FIGNL1 complex control RAD51 dynamics at the site of DNA breaks. In line with this model, clustering of TurboID preys based on their chemogenomic profiles^28^ revealed that C1orf112 and FIGNL1 cluster with RAD51 paralog XRCC3 (**Fig 5D, Table S4**). To further test this model, we investigated whether C1orf112 impacts the dynamic of RAD51 foci formation and resolution in cells treated with an ICL-inducing agent. RAD51 foci rapidly appeared 3-6 hours after the addition of cisplatin to the media (**Fig 5E and S5G**). While more foci accumulated in cells depleted for C1orf112, their intensity were like the intensity of the foci in cells treated with a siRNA control (**Fig 5F**), suggesting that RAD51 is properly loaded at ICLs in these cells. In contrast, more than 40% of C1orf112-depleted cells still exhibited more than 5 RAD51 foci 72 hours post-treatment, a time frame that was sufficient to resolve these foci in the control condition. Altogether, these findings demonstrate that C1orf112 regulates RAD51 unloading at ICLs following strand invasion by regulating its interaction with FIGNL1 (**Fig 5G**).

## DISCUSSION

### C1orf112 is a novel regulator of HR-mediated ICL-repair

While the initial steps involved in ICL repair have been extensively described, it remains largely unclear how ICL intermediates are processed and subsequently resolved. CRISPR-based functional genomics have been a powerful strategy to gain novel insight into several DNA repair pathways^28^, including the recent characterization of SCAI as a modulator of repair pathway choice during ICL processing^7,8^. In this work, we delineated the contribution of the poorly characterized C1orf112 in the latter stage of ICL repair using a similar strategy. We show that lack of C1orf112 impairs cell survival in response to a series of ICL-inducing agents. In these conditions, C1orf112 acts downstream of the FA pathway where it is functionally epistatic with the AAA+ ATPase FIGNL1 for the resolution of HR-mediated intermediate. Specifically, our findings reveal that C1orf112 is required for the resolution of RAD51, a phenotype that is reminiscent of the RAD51 unloading activity of FIGNL1^9^. Consistent with a coordinate role of C1orf112 and FIGNL1 in regulating HR-mediated DNA repair, mutation in *figl1-1*, the ortholog of FIGNL1 in plants, is epistatic with *flip-1* mutation, the ortholog of C1orf112, in limiting meiotic crossover formation^35^. While we were unable to detect the recruitment of C1orf112 at ICLs, previous studies identified this factor at stalled replication fork by iPOND^36,37^, suggesting that they specifically contribute to ICL-induced replication fork collapse^1^. The fact that C1orf112 limits the formation of micronuclei, the accumulation of 53BP1 nuclear bodies, and prevents the induction of senescence, supports a model where C1orf112 may act in the response to replication stress.

### C1orf112 forms a heterodimer with FIGNL1 to regulate RAD51 accumulation upon repair

Previous reports identified FIGNL1 as key regulator of the resolution of RAD51 foci in different biological contexts^9,17,32^. Interestingly, our systematic proteomics analysis of C1orf112 identified FIGNL1 as a constitutive proximal interactor that we validated *in cellulo*. Our preliminary data suggest that this heterodimer is necessary for the stability of the complex (**Fig. S5I**), reminiscent of what has been previously observed with the BRCA1/BARD1 heterodimer, where each member of the heterodimer controls the abundance, stability, and function of the other^38,39^. Our findings suggest that C1orf112 acts as a scaffolding platform for FIGNL1 in complex with RAD51, thereby regulating its unloading. Furthermore, our proximal mapping of FIGNL1 identified the RAD51 paralog XRCC3, a component of the CX3 complex, which has been suggested to act downstream of RAD51 loading^40^. Although RAD51 was recently involved in the protection of cells from transcription-replication conflicts^41^, either C1orf112- or FIGNL1-depleted cells are not sensitized to transcription-interfering agents such as Trabectedin and illudinS (**Fig S6**), suggesting that the heterodimer specifically regulates RAD51 dynamics following strand invasion and not at stalled replication forks per se. Interestingly, cells depleted from either one of these factors are specifically sensitive to chemical agents that generate replication roadblocks, which requires a specific processing before repair such as ICL-agents and agents that promote either base oxidation or base alkylation (**Fig S6**)^42–44^ Of note, our data suggest that C1orf112 and the SWS1-SWSAP1-SPIDR complex, which inhibit FIGNL1-dependent RAD51 unloading^9,10^, play opposite roles in response to replication stress. Consistently, both complexes cluster differently when analysed by chemogenomic profiling (**Fig S6**).

### FIGNL1-independent function of C1orf112

Chemogenomic profiling of both FIGNL1 and C1orf112 highlights the differential contribution of each factor to cell survival upon genotoxic stress, suggesting that C1orf112 and FIGNL1 also have distinct functions in cells. Consistent with this hypothesis, FIGNL1 is highly conserved in a range of eukaryotic species while C1orf112 is not conserved in Fungi as well as in model organisms such as C. elegans and D. melanogaster^32^. In line with this observation, the predicted C1orf112 interacting region is located in a less conserved region of FIGNL1. Alike other scaffolding proteins such as REV7, it is possible that C1orf112 forms different molecular complexes, depending on the cellular context, thereby preserving genomic stability (reviewed in^45^). In fact, our proteomic analysis also identified ERCC6L as one of the top constitutive interactors of C1orf112, which may likely reflect a role during mitotic progression^18^.

### Clinical relevance of C1orf112 in cancers

So far, no mutation in C1orf112 has been linked to a FA-related phenotype. However, high levels of C1orf112 have been associated with poor outcomes in a series of cancers, including gliomas^46^, and therefore may be a potential prognostic marker in multiple tumor types^47^. Our data further suggest that C1orf112 expression may modulate the response to different well-established chemotherapeutic agents, thereby influencing patient outcome.

## METHODS

### Cell Culture and plasmids transfections

All cell lines were maintained at 37°C and 5% CO2. Namalwa and Raji Burkitt lymphoma cell lines (kind gift of Dr. Jerry Pelletier, McGill university) were cultured in Roswell Park Memorial Institute (RPMI) 1640 medium (Wisent) and supplemented with 20% fetal bovine serum (FBS, Sigma) and 1% Penicillin-Streptomycin (Wisent). RPE1-hTERT p53 WT and p53^-/-^ cells (kind gift of Dr. Daniel Durocher, University of Toronto) and IMR90 cells were cultured in Dulbecco’s modified Eagle medium (DMEM) medium (Wisent) and supplemented with 10% FBS and 1% Penicillin-Streptomycin. U2OS (purchased from ATCC) and U2OS 2-6-5 cell lines (a kind gift from Roger Greenberg (University of Pennsylvania, Philadelphia, PA)^24^) were cultured in McCoy’s medium (Life Technologies) and supplemented with 10% fetal bovine serum (FBS). Hela DR-GFP cells, U2OS DR-GFP and U2OS SA-GFP cells (kind gift of Dr. Jeremy Stark, City of Hope) were cultured in DMEM and McCoy’s medium (Wisent), respectively. These cells were supplemented with 10% FBS and 1% Penicillin-Streptomycin.

RPE1-hTERT Cas9 p53^-/-^ C1orf112^-/-^ cells were generated by transduction of the parental cell line with lentiviral vectors harboring C1orf112-sgRNAs targeting exon 5 and exon 13 (**Table S5**). Cells were selected with puromycin for 4 days, and single clones isolated by single-cell sorting. Knockout cells were validated using immunoblotting. Stable Cas9-expressing Burkitt lymphoma cell lines (Namalwa and Raji Burkitt cells) were generated using the LentiCas9-Blast vector as previously described^48^ and validated for Cas9 expression by immunoblotting analysis. Cell lines used for all mturboID-MS experiments were generated in HEK293 Flp-In^TM^ T-REx^TM^ cells as previously described^14^ and pool of stable transfectants selected with 200 μg/mL hygromycin (Multicell) and 5 μg/mL blasticidin (Invivogen). Expression of bait proteins were induced with 1 μg/mL tetracycline for 24 h. Plasmid transfections were done using Lipofectamine™ 2000 (Invitrogen) and TransIT-LT1 (Mirus) transfection agents according to the manufacturer’s instructions. All cell lines were validated using short tandem repeat (STR) markers and tested negative for mycoplasma contamination.

### RNA interference

The pLKO shRNA plasmids against C1orf112 were obtained from the McGill Platform for Cellular Perturbation (MPCP) as part of the TRC/RNAi Consortium from the Broad institute. Nontargeting shRNA control was purchased from Addgene. Single siRNA duplex sequences targeting C1orf112, FIGNL1, FANCA, a custom non-targeting sequence, and SMARTPool siRNAs targeting RAD51 or CtIP were purchased from Dharmacon. Unless stated otherwise, all siRNAs were transfected at a concentration of 25 nM in a forward transfection for 48 hours using RNAimax (Invitrogen) according to the manufacturer’s instructions. Knockdowns were confirmed by immunoblotting or qPCR analyses. All siRNA and shRNA sequences are detailed in **Table S5**.

### RNA extraction and RT-qPCR

Total RNAs were extracted using the RNeasy mini kit according to the manufacturer’s instruction (QIAGEN) and quantified using a NanoDrop™ spectrometer. 500 ng of total RNA was reversed transcribed with the High-Capacity cDNA reverse transcription (RT) kit (Invitrogen) in accordance with the manufacturer’s instructions. Prior to the RT, contaminant genomic DNA was removed by DNaseI (ThermoFisher) treatment and confirmed by GAPDH RT-PCR on DNaseI-treated reactions. qPCRs were performed on a LightCycler 480 apparatus (Roche) with the LightCycler 480 SYBR Green 1 qPCR master mix (Roche) using the following program: 40 cycles of 94°C denaturation for 15 seconds, 56°C annealing for 5 seconds and 72°C elongation for 15 seconds. 5% of the RT-PCR reaction was used as a template. The standard curve was calculated with serial dilution of the U2OS cDNA as templated and used to determine the relative expression of each gene before normalization to the relative expression of GAPDH. The PCR primers are listed in **Table S5**.

### Chemicals and sources of DNA damage

In the FOK1 system, DSBs were induced at the *LacO* array by inducing the nuclear localization (4-Hydroxytamoxifen (4-OHT, 100 nM, #3412, Tocris)) and stabilization (Shield-1 ligand, 0.5 μM, CIP-S1-0001, CheminPharma)) of the ER-mCherry-LacR-FOK1-DD nuclease for 4 hrs prior to immunofluorescence sample preparation. DNA damages were induced by exposing cells to either ionizing irradiation (IR 1 or 10 Gy) with a CellRad (Precision X-Ray Inc.), cisplatin treatments (Tocris 1 μM for 3 or 24 hrs, 250 nM in time course experiment, and 2 μM in survival assay), mafosfamide (MAF, Toronto Research chemicals, 1.30 μM and 1.57 μM in Namalwa and Raji screens respectively), neocarzinostatin (NCS) (Sigma-Aldrich, 100 ng/mL), hydroxyurea (HU, Sigma-Aldrich, 4 mM and 1 mM in survival assay), formaldehyde (Sigma-Aldrich, 150 μM), mitomycin C (MMC, Sigma-Aldrich, 150 nM), Talazoparib (BMN 673, Selleck Chemicals, 3 μM), or UV (10 J/m^2^). UV irradiations were carried on with a germicidal lamp (243 nm).

### Cell viability assays

U2OS cells were seeded in 6-well plate (75 000 cells/wells) 24 hrs prior to siRNA transfection (siCtrl, siC1orf112 or siFIGNL1). Forty-eight hrs post-transfection, cells were exposed to the indicated treatment. Following a recovery of 48 hrs, cells were harvested and counted using an hematocyter

### Plasmids

DNA sequences of sgRNAs were cloned into a modified form of LentiGuide-puro with BFP or mCherry marker as described in^49^. The cDNA for C1orf112 (MHS1010-202739778, *Horizon Discovery*) and FIGNL1 (MHS6278-202759761, *Horizon Discovery*) were obtained from Dharmacon, and coding sequences with respective attB1/B2 adapters were PCR-amplified and subcloned into pDONR221 vectors using a BP clonase II reaction according to the manufacturer’s instructions (Invitrogen), to generate ENTRY vectors. ENTRY clones containing C1orf112 constructs (aa 1-175, aa 1-735) and FIGNL1 constructs (aa 1-120, aa 121-674, aa 1-360, aa 361-674, aa 295-344) were generated using the same approach. FIGNL1 mutations and depletion construct (Δ295-344, F295E, K447A, D500A and K477A/D500A) were directly introduced in ENTRY vectors using the Quick-change approach. Quick-change Site-Directed Mutagenesis was conducted using primers listed in **Table S4** using the Quick-change (Agilent) or Q5-Site-directed mutagenesis kit (NEB) following the manufacturer’s protocol. LR recombining reactions were then conducted to transfer ENTRY constructs into pDEST-mturboID (kind gift from Dr. Anne-Claude Gingras, University of Toronto), pDEST-mCherry-LacRnls (kind gift from Dr. Daniel Durocher, University of Toronto), or pDEST-FRT-TO-eGFP-nls. All sequences were validated through Sanger sequencing. A list of plasmids used in this study is provided in **Table S7**.

### CRISPR-Cas9-based genome-wide screening

Namalwa and Raji genome-wide screens were executed as previously described^50^. Briefly, 245 millions Namalwa and Raji cells expressing Cas9 were transduced with TKOv1 sgRNA lentiviral library at MOI (0.2), ensuring coverage of at least 500-fold for each individual sgRNA represented in the cell population. Two days later, the selection of fully edited transduced cells was achieved by adding puromycin to the media at a final concentration of 2 μg/ml for 7 days. Following a 2-day recovery in fresh media without selection, cells were split into 2 pools, in triplicate, at a cell density of 45 million cells (D0). The first pools were treated with mafosfamide at its IC25 (1.30 μM and 1.57 μM in Namalwa and Raji screens respectively) and the second pool with DMSO as a vehicle. Cells were then cultured for 14 doubling times with puromycin at a concentration of 1 μg/ml. Cells were counted and passaged every 3 days at a cell density of 0.37 million cells per mL to maintain fold coverage of 500 cells per sgRNA, until D14. At each time point, cell pellets were frozen for subsequent genomic DNA purification. Genomic DNA isolation was performed as described in^14^ For next-generation sequencing (NGS) library preparation, sgRNAs were amplified from genomic DNA using two rounds of nested PCR with inner oligo and outer oligo annealing. The initial outer PCR was performed with TaKaRa ExTaq^®^ DNA Polymerase Hot-Start Version polymerase (Takara) using forward (FW) and reverse (RV) outer primers (**Table S5**). PCR products were pooled, and ~2% of the input was amplified using Hot start Q5 polymerase to add Illumina HiSeq adapter sequences (**Table S5**). The resulting product from each pooled sample was further purified following separation on an 8% 0.5× TBE polyacrylamide gel. The library NGS was quantified using qPCR and analyzed by deep-sequencing on the HiSeq 2500 Illumina platform. Reads were trimmed of NGS adapter sequences using the Cutadapt tool. Reads were aligned to the sgRNA library index file using Bowtie2. BAM files were generated using samtools, and total read count tables were subsequently generated using the MAGeCK count command. DrugZ algorithm was used to identify gene knockouts which were depleted or enriched from D14 populations in comparison to D0^16^.

### Gene Set Enrichment Analysis (GSEA) and Network interaction analysis

GSEAs were conducted individually for each DrugZ-ranked list from the Namalwa and Raji screens using the GSEA v4.3.0 package and using the Gene Ontology biological process gene sets. For the integrated pathway enrichment analysis, an average of the NormZ values acquired individually for both screens using DrugZ was obtained, and a merged ranked list was generated. Pathway enrichment analysis on the integrated ranked list was conducted using the package Enrichr as previously described^51,52^. Network interaction analysis was performed using Cytoscape v 3.9.1 and the GeneMANIA package^53,54^

### Construction of heatmaps

All heatmaps were generated with the ComplexHeatmap R package^57^ by performing hierarchical clustering on drugs NormZ-scores derived from published CRISPR screens^28^ and limited to preys identified by our miniturboID assays. In legends, dark blue indicates NormZ scores ≤ −3 while bright red indicates scores ≥ 3. Similarly, we created a heatmap out of our miniturboID results using the log2-transformed SAF (Spectral Abundance Factor) metric, a normalization method calculated by dividing average spectral counts of preys by their respective protein length in amino acids.

### Acquisition of immunoblotting images

All immunoblot images were acquired with an Azure Biosystems c300 Imaging System instrument. Briefly, membranes were exposed to Azure Radiance ECL (VWR) for 1 min before acquisition on the Azure apparatus with the sensitivity parameter set to “Normal” (1108×834). Whole pictures were adjusted for luminosity and contrast with the Adobe Photoshop software.

### CRISPR-based competition growth assay

Growth competition assays were conducted as described in^19^ and^49^. Briefly, RPE1-hTERT Cas9 p53^-/-^ were transduced with lentiviral particles of Lenti-mCherry-sgRNA-LacZ or Lenti-BFP-sgRNA-GOI (C1orf112/FIGNL1/FANCA/MRE11). Twenty-four hours post-transduction, cells were selected with 15 μg/mL of puromycin for 4 days. BFP and mCherry labelled cells were mixed at a 1:1 ratio and seeded in a 6-well plate. At the initial time point (T0), cells were treated with cisplatin at 6 μM (IC25) or vehicle (DMSO), and the levels of each fluorophore were measured via flow cytometry. Cells were maintained under these conditions and subcultured for 20 days.

The ratio of BFP to mCherry fluorescent cell population were assessed via flow cytometry every 5 days.

### Apoptosis

RPE1-hTERT Cas9 p53^-/-^ C1orf112^-/-^ clones 1 and 2 were seeded in 6-well plates and treated with an overnight treatment of 4 μM cisplatin. The next day, cells were washed with Dulbeccos’s PBS and fresh media added to the plate. Forty-eight hours post cisplatin treatment, cells were processed for Annexin-V (Biolegend) and PI staining, following the manufacturer’s instructions. Analyses of Annexin-V and PI signals were done on at least 30 000 events acquired on a BD Fortessa (Becton Dickinson). Data were analyzed using the FlowJo software as previously described^55^.

### Senescence-associated β-Galactosidase Assay

Senescence-associated β-galactosidase (SA β-gal) assays were performed as previously described^56^. Briefly, shRNA expressing REP1 hTERT p53WT or p53^-/-^ or IMR90 cells were fixed with 0.5% glutaraldehyde in PBS, washed and kept at 4 °C in PBS supplemented with 1 mM MgCl2 (pH 6). Cells were stained with X-Gal solution containing potassium ferricyanide in PBS supplemented with 1 mM of MgCl2 (pH 6). Images were acquired, and the percentage of SA β-gal positive cells was quantified.

### GFP-based DNA repair Assays

DNA repair by HR (DR-GFP) or SSA (SA-GFP) of I-SceI-generated DSB were measured as previously described^14,26^. Briefly, HeLa DR-GFP, U2OS DR-GFP or U2OS SA-GFP cells were seeded in 6-well plates at a density of 100,000 cells/well and were transfected with 25 nM of siRNA. Twenty-four hours later, cells were transfected with pCBA-SceI plasmid using Lipofectamine 2000 (Invitrogen). Cells were harvested 48 hrs post-transfection, and the percentage of GFP-expressing cells was measured by flow cytometry. Analysis of GFP-positive signal was done on at least 30 000 events acquired on a BD Fortessa (Becton Dickinson). Data were analyzed using the FlowJo software and presented as previously described^14^

### Expression profiling IC50

The correlation between gene expression and drug sensitivity was conducted using expression datasets for each gene of interest available from the Cancer Cell Line Encyclopedia (CCLE) project as part of the Cancer Dependency Map (DepMap). IC50 values were obtained from the Genomics of Drug Sensitivity in Cancer (GDSC) database (Release 8.4, GDSC2 datasets)^58^.

### Sulforhodamine B (SRB) Assay

RPE1-hTERT cells were seeded in 96-well plates at a density of 1000/cells per well. Twenty-four hours later, cisplatin and mafosfamide were added in a two-fold serial dilution from 50 to 0.097 μM. Survival was assessed four days after treatment using the sulforhodamine B (SRB) colorimetric assay as described previously^59,60^. Briefly, after drug treatment cells were fixed by adding 100 μL of 10% trichloroacetic acid (TCA, Bioshop Canada) and incubated at 4°C for 1 hr with gentle agitation. Cells were washed four times and plates were left air-drying overnight at room temperature. The next day, cells were stained with 100 μL of 0.057% SRB (Sigma Aldrich) and incubated at room temperature for 30 minutes. Plates were then rinsed four times using 1% acetic acid and were left air-drying overnight. Protein content was solubilized by adding 200 μL of 10 mM Tris base solution (pH 10.5) for 30 minutes at room temperature. Measurement of optical density (OD) at 510 nm was conducted using a FLUOstar Optima microplate reader. Background correction was conducted using the measurement of control wells with media. Treatments were performed in triplicate, averaged, and normalized to untreated control. IC50 concentrations were obtained using the slope’s equation for log(concentration of drug) vs normalized OD.

### Immunofluorescence Microscopy

U2OS and U2OS 2-6-5 cells were grown on glass coverslips in 24-well plates and fixed with 2% (wt/vol) paraformaldehyde (PFA) in PBS for 20 minutes at RT. Exceptionally, cells used in the time course experiments were fixed with 100% MeOH for 20 minutes at −20°C. When indicated, PFA-fixed cells were processed for immunofluorescence by further permeabilizing cells with 0.3% (vol/vol) Triton X-100 for 20 minutes at RT. Cells were then incubated with blocking buffer (10% goat serum, 0.5% NP-40, 0.5% saponin in 1X PBS) for 30 minutes at RT and then incubated with primary antibodies (**Table S6**) in blocking buffer for 2 hrs at RT. After three washes with PBS, cells were incubated for 1 hr at RT with secondary antibodies (**Table S6**) and 4’,6-diamidino-2-phenylindole (DAPI, 0,4 μg/mL) blocking buffer. Coverslips were mounted onto glass slides with ProLong™ Diamond Antifade Mountant agent (Invitrogen). Images were acquired using a Zeiss LSM900 laser-scanning microscope equipped with a 48X and 63X oil lens. In all micrographs, dashed lines indicate the nucleus outline, and insets represent a 10-fold magnification of the indicated fields. Each quantification was done on at least 3 biological replicates, and at least 100 cells or 50 foci were counted in each experiment. Unless stated otherwise, significance was assessed by performing a t-test with Welch’s correction.

### mturboID sample preparation for mass spectrometry

HEK293 Flp-In^tm^ cells expressing mTurboID-tagged protein or mTurbo tag alone were seeded in 150 mm plates in technical duplicates. Induction of fusion protein and proximity biotinylation were conducted as previously described^14,55^. Briefly, induction of mturboID-tagged protein expression was done by adding 1 μM tetracycline to the media for 24 h. Then, the medium was supplemented with 50 μM biotin for 1 hr and incubated for an additional 3hrs with 100 ng/mL neocarzinostatin (NCS). Cells were harvest, washed and lysed in ice-cold RIPA buffer (50 mM Tris-HCl pH 7.4, 150 mM NaCl, 1% NP-40, 1 mM EDTA, 0.1% SDS and 0.5% sodium deoxycholate, 1 mM PMSF, 1 mM DTT) supplemented with 1:500 Sigma-Aldrich protease inhibitor cocktail P8340. and sonicated on ice. After sonication, 250 units of benzonase (EMD) were added sample prior to a 30 minutes centrifugation at 12,000 xg at 4°C. Supernatants were transferred to pre-washed streptavidin-Sepharose beads (GE, #17-5113-01) and incubated at 4°C on a rotator for 3 hrs. Beads were centrifuged at 400 xg for 1 minute and sequentially washed 2 times with RIPA buffer and 3 times with of 50 mM ammonium bicarbonate (ABC, pH 8.0). Beads were then resuspended in 100 μL of ABC, and on-bead digestion was achieved by adding 10 μg of trypsin (Sigma) to the suspension for overnight digestion at 37°C. The next day, an additional 10 μg of trypsin was added to each sample, and further digested for 3 hrs. Beads were pelleted by centrifugation for 1 min at 400 g, and the supernatant-containing peptides were pooled with supernatants from two subsequent rinses with 100 μL of mass spectrometry grade H_2_O. Formic acid was added to the pooled samples to a final concentration of 5% to end digestion. Samples were centrifuged at 16,000 xg for 10 minutes at room temperature and supernatants were lyophilized using vacuum centrifugation. Dried peptides were kept at −80°C.

Analysis of MS data was conducted as described before with minor modifications^55^. Briefly, samples were injected into an Orbitrap Fusion (Thermo Fisher), and raw files were analyzed with the Comet, XTandem! and Mascot search engines using the human RefSeq database (version 20170518) supplemented with ‘‘common contaminants’’ from the Max Planck Institute (http://maxquant.org/downloads.htm), the Global Proteome Machine (GPM; http://www.thegpm.org/crap/index.html) and decoy sequences. The search parameters were set with trypsin specificity (two missed cleavage sites allowed), variable modifications involved Oxidation (M) and Deamidation (NQ). The mass tolerances for precursor and fragment ions were set to 10 ppm and 0.6 Da, respectively, and peptide charges of +2, +3, +4 were considered. Search results were individually processed by PeptideProphet^61^, and peptides were assembled into proteins using parsimony rules first described in ProteinProphet^62^ using the Trans-Proteomic Pipeline (TPP). TPP settings were the following: -p 0.05 -x20 -PPM –d “DECOY”, iprophet options: pPRIME and PeptideProphet: pP. To estimate the interactions statistics, we used SAINTexpress (PMID 24513533) (version 3.6.1) on proteins with iProphet protein probability ≥ 0.9 and unique peptides ≥ 2. Each bait was compared against its respective negative control (treated or not with NCS). These controls comprised of pulldowns from HEK cells expressing the empty vector (without the BirA*-Flag) in triple technical replicates. SAINT analyses were performed with the following compression settings: nControl:2, nCompressBaits:2. Interactions displaying an average probability (AvgP) ≥ 0.7 were considered as statistically significant (Supplementary Table X). Unfiltered contaminants, such as Keratin, BirA* and beta galactosidase were discarded in every bioinformatics analyses. Using the ProHits-Viz online tool (prohits-viz.org), we generated a dot plot representing the relative AvgSpec of identified proteins for both baits.

## ACKNOWLEDGMENTS

We are grateful to members of the Orthwein and Fradet-Turcotte laboratories for critical reading of the manuscript; Daniel Durocher, Roger Greenberg and Jeremy Stark for plasmids and other reagents. EPC is a recipient of a doctoral fellowship from the Fonds de Recherche Santé Québec (FRQS). Work in the A.O. laboratory was supported by a CCSRI Innovation Grant (#706389), a Transition Grant from the Cole Foundation and an internal Operating Fund from the Sir Mortimer B. Davis Foundation of the Jewish General Hospital. Work in the A.F.-T laboratory was supported by a Canadian Institutes of Health Research Grant (PDT_152948 to A.F.-T.). Work in the J.-F.C. laboratory is supported by a Canadian Institutes of Health Research Grant (PJT-178241). A.F.-T. is a Tier 2 Canada Research Chair in Molecular Virology and Genomic Instability and is supported by the Foundation J.-Louis Lévesque. J.-F.C. holds the Tier 1 Canada Research Chair in Cellular Signalling and Cancer Metastasis and is supported by the Alain Fontaine Chair in Cancer Research from the IRCM Foundation.

## AUTHOR CONTRIBUTIONS

E.P.C., J.D., F.M., A.G., A.O. and A.F-T. designed research; E.P.C., J.D., R.V., L.S., A.M., J.B., E.G.L., V.L., A-M.L., and J-F.C. performed research and analyzed data; A.F.-T. and A.O. wrote the original draft and E.P.D. and J.D. edited the manuscript.

## CONFLICT OF INTEREST

The authors declare that they have no conflict of interest.

## SUPPLEMENTARY FIGURE LEGENDS

**Figure S1.**
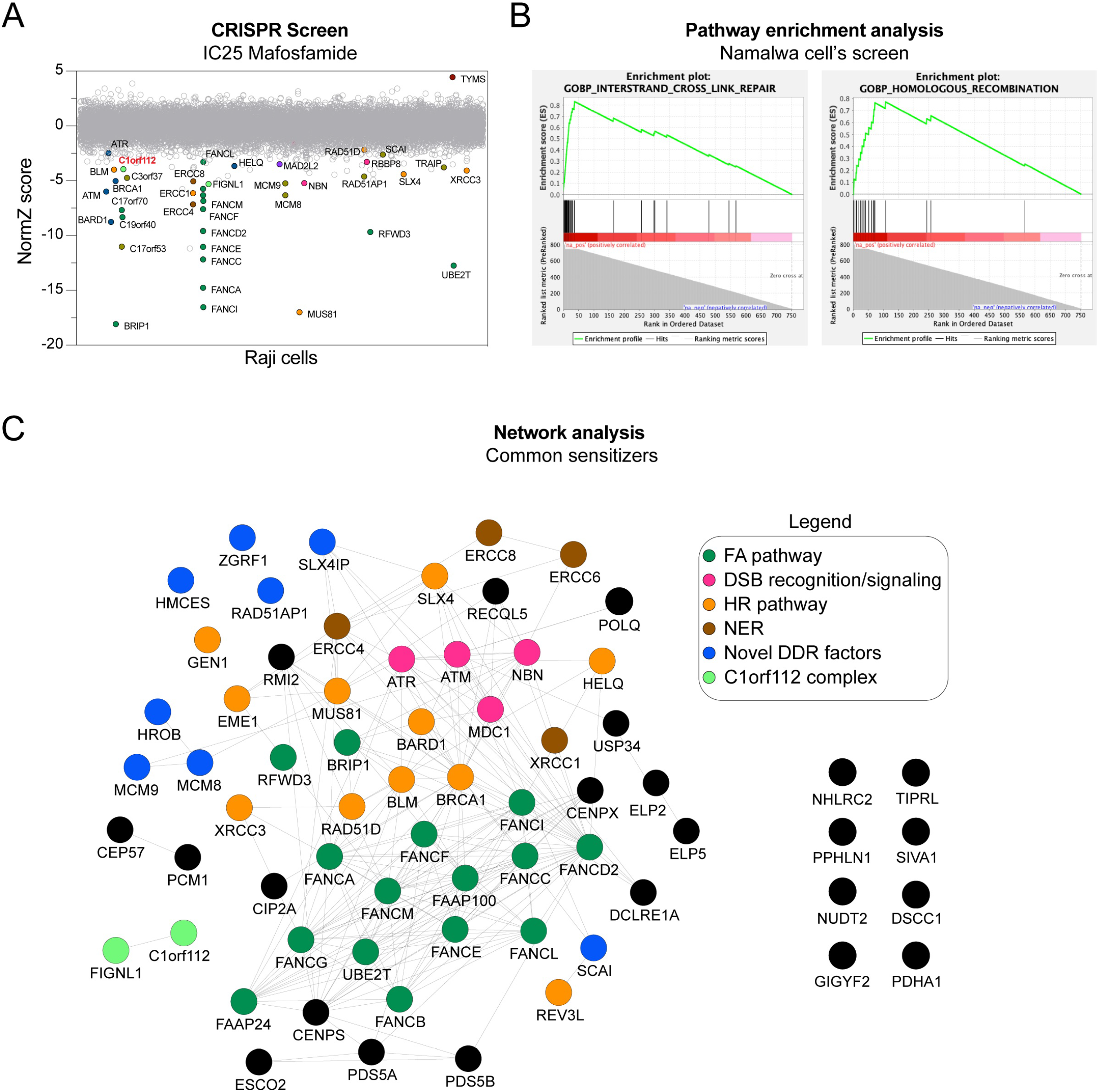
Additional analysis of CRISP screen performed in Raji and Namalwa cells, related to Fig. 1. (A) Horizontal scatter plot of DrugZ-generated ranking of the Raji mafosfamide CRISPR screen. NormZ values are plotted on the Y-axis, and gene names are plotted on the X-axis. (B) Gene Set Enrichment Analysis (GSEA) analysis of the score obtained in the Namalwa screen for ICL repair and HR. (C) Network analysis displaying protein physical interactions for sensitizer genes common to Namalwa and Raji CRISPR screens using Cytoscape and the GeneMANIA package. (D) WCE of RPE1-hTERT WT or p53^-/-^ clonal cells depleted or not for C1orf112 with the indicated sgRNA were analyzed by immunoblot with an anti-C1orf112 antibody. Anti-β-actin was used as a loading control. (E) Competitive growth results obtained in RPE1-hTERT p53^-/-^ cells targeted with a sgRNA against C1orf112. Cells were either treated with DMSO as a vehicle or cisplatin IC25 = 2.4 μM. Data are represented as the mean ± SEM (n = 3 technical replicates).

**Figure S2.**
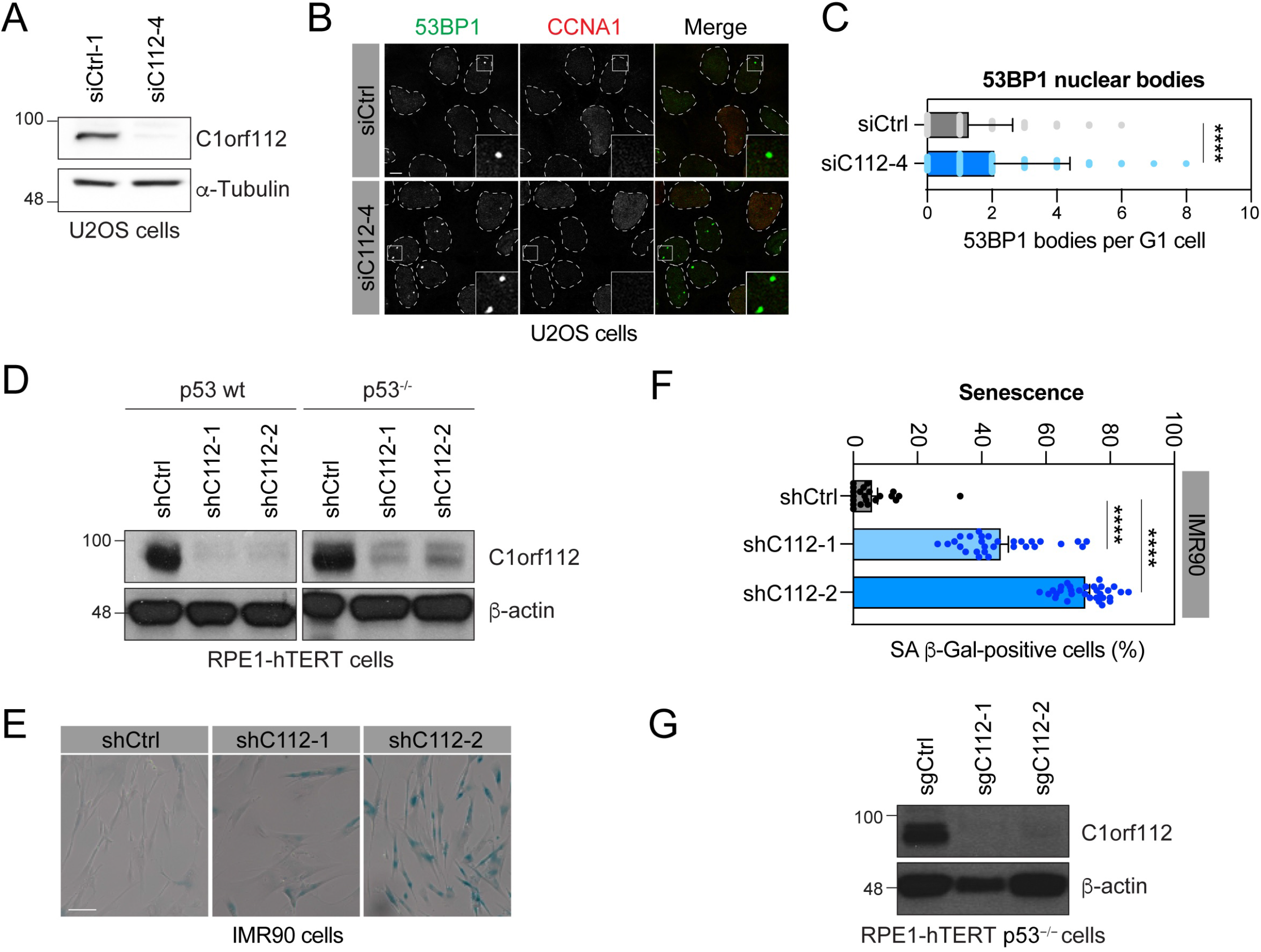
Validation of C1orf112 knockdowns and knockouts, 53BP1 nuclear bodies in analysis and additional senescence assay in IMR90 cells, related to Fig.2. (A) WCE of U2OS cells depleted or not for C1orf112 were analyzed by immunoblot with an anti-C1orf112 antibody. Anti-a-Tubulin was used as a loading control. (B) Representative images of U2OS cells treated with a siC1orf112 or non-targeting siRNA for 48 hrs were processed for 53BP1 and CCNA1 (Cyclin A) immunofluorescence. Scale bar = 5 μm. (C) Quantification of 53BP1 nuclear bodies per CCNA1-negative cell as shown in (B). Data are represented as the mean ± SD (n = 3 independent experiments. A minimum of 100 cells were analyzed per condition per experiment). Data were analyzed with an unpaired t-test with Welsh’s correction. (D) WCE of RPE1-hTERT WT or p53^-/-^ cells depleted or not for C1orf112 with the indicated shRNA were analyzed by immunoblot with an anti-C1orf112 antibody. Anti-β-actin was used as a loading control. (E) Representative images of IMR90 cells depleted or not for C1orf112 and stained with SA β-galactosidase. Scale bar = 5 μM (F). Quantification of SA-β-gal positive cells as shown in (E). Data are represented as the mean ± SEM (n = 3 independent experiments). Data were analyzed with an ordinary one-way ANOVA test with Dunnett’s multiple comparison tests. ****p<0.0001

**Figure S3.**
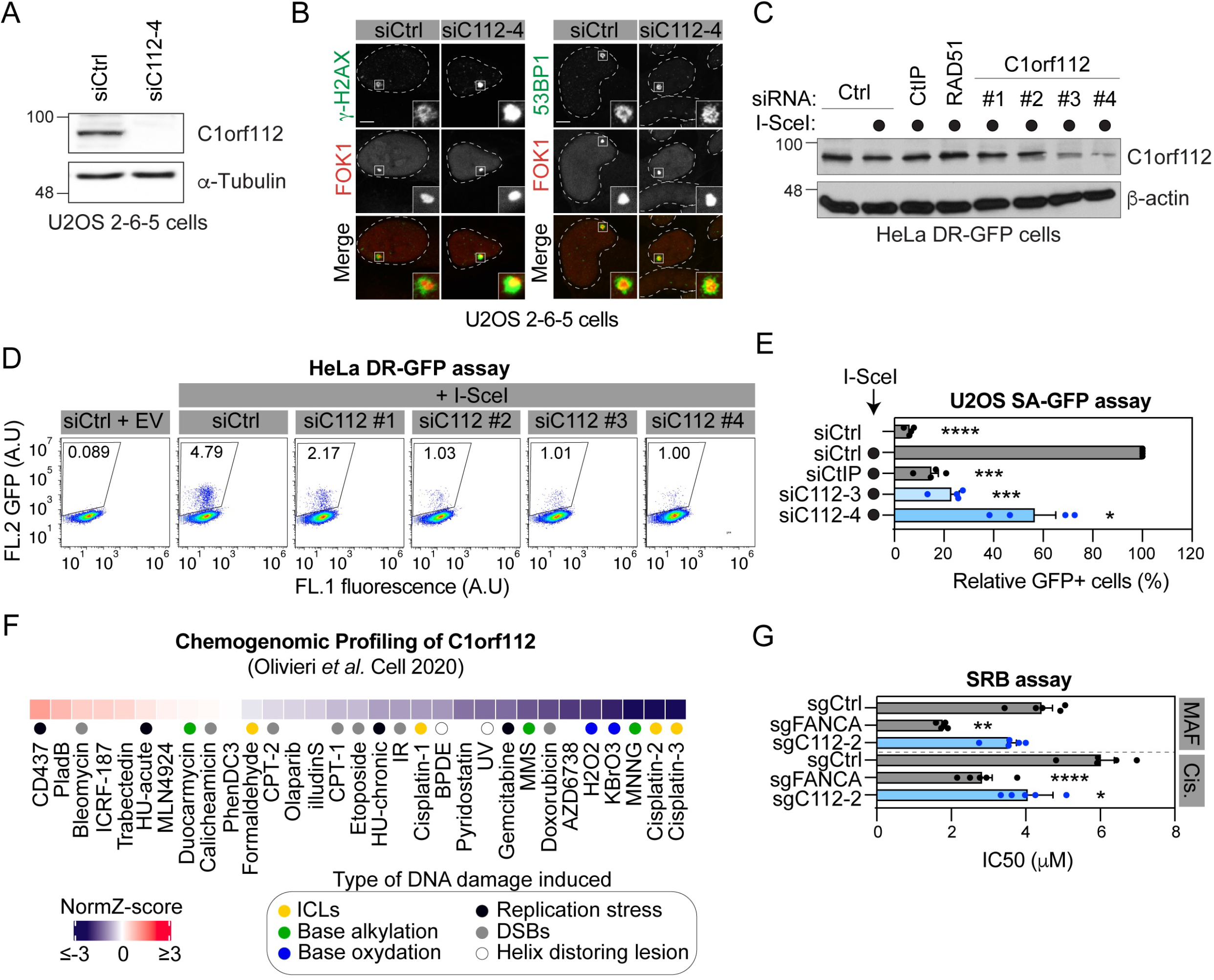
Characterization of C1orf112 knockdown and knockout cells related to Fig 3. (A) WCE of U2OS 2-6-5 cells depleted or not for C1orf112 were analyzed by immunoblot with an anti-C1orf112 antibody. Anti-a-Tubulin was used as a loading control. (B) Representative images of U2OS 2-6-5 cells depleted or not for C1orf112. Cells were treated as in Figure 3E and processed for γ-H2AX (left) and 53BP1 (right). Scale bar = 5 μm. (C) WCE of HeLa DR-GFP depleted or not for CtIP, RAD51 or C1orf112 were analyzed by immunoblot with an anti-C1orf112 antibody. Anti-β-actin was used as a loading control. (D) Representative scatter plots of GFP-positive HeLa DR-GFP cells analyzed by flow cytometry and presented in Figure 3G. Flow cytometric profiles where green fluorescence (FL2) and auto orange fluorescence (FL1) are plotted on the y-axis and x-axis, respectively. (E) Quantification of GFP-positive U2OS SA-GFP cells depleted or not for CtIP or C1orf112 Twenty-four hours post siRNA transfection, cells were transfected with the I-SceI expression plasmid (●) or an empty vector. Data are represented as the mean ± SEM (n = 4 independent experiments, 30,000 cells quantified per experiment in each condition). Data were analyzed using a one-way Welch’s ANOVA test and Dunnett’s multiple comparison tests. (F) Chemogenomic profiling using the NormZ-score from the CRISPR screens performed in RPE1 hTERT Cas9 p53^-/-^ cells with the indicated DNA damaging agents as detailed in^28^. (G) Quantification of RPE1-hTERT p53^-/-^ cells survival in response to ICL-inducing agent. Cells were transduced with lentiviral particles containing sgRNAs against C1orf112, FANCA or a non-targeting control (sgCtrl). After selection with puromycin, cells were seeded in 96-well plates and treated with mafosfamide (MAF) or cisplatin (Cis.). Both drugs were added at a maximum concentration of 50 μM in a two-fold serial dilution until 0.097 μM. Data are represented as the mean ± SEM (n = 5 independent experiments). Data were analyzed using a one-way Welch’s ANOVA test and Dunnett’s multiple comparison tests. *p<0.05, **p<0.01***p<0.001, ****p<0.0001

**Figure S4.**
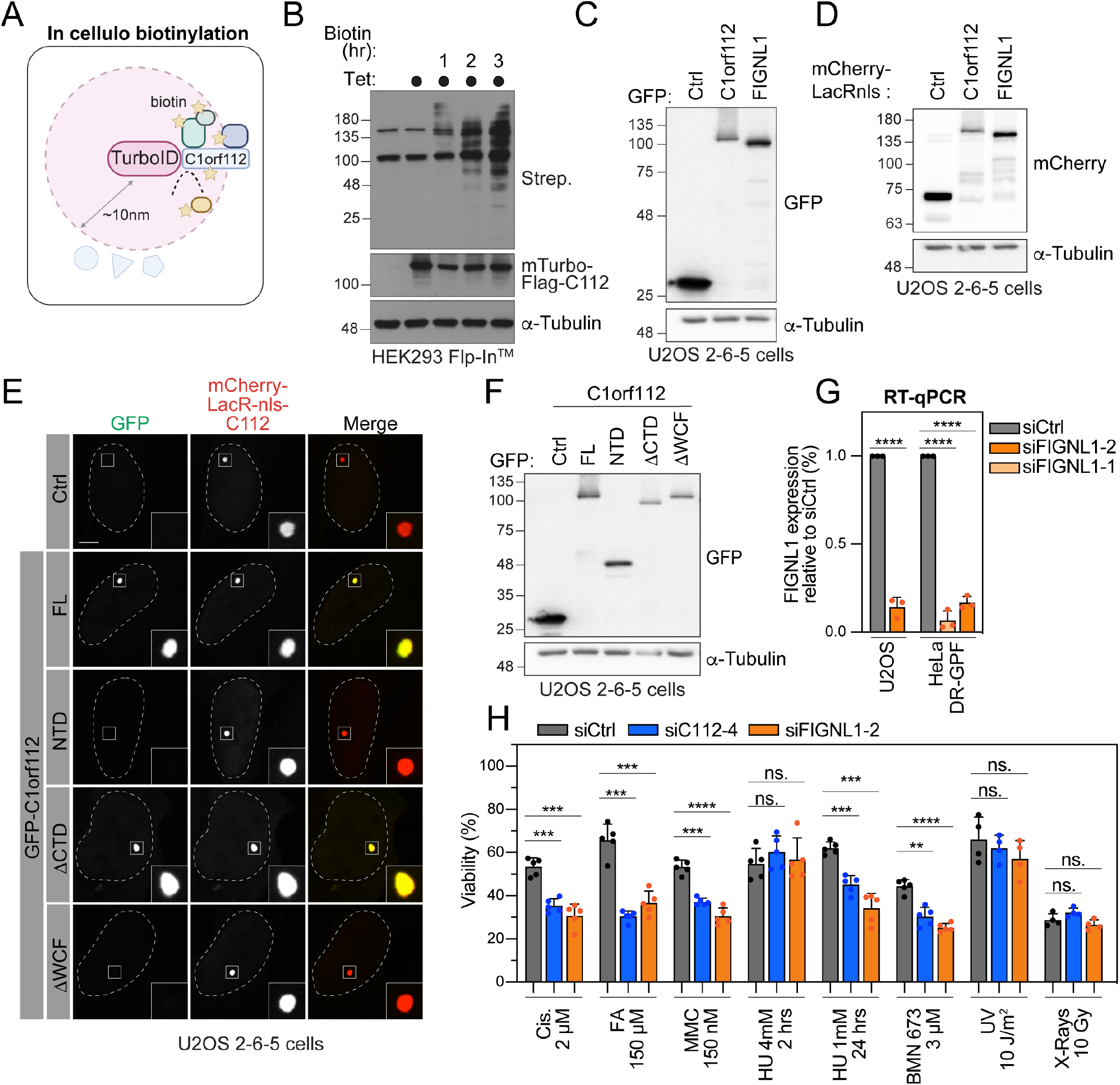
Validation of protein expression and cell lines for mTurboID and LacO/LacR analysis, and characterization of C1orf112 and FIGNL1 in survival assays and patients. (A) Schematic representation of the biotinylation of C1orf112 proximal endogenous proteins using TurboID technology. Cutoff for proximity tagging is reported at 10nm^29^. (B) WCE extracts of HEK293T FL/IN cells induced or not for mturbo expression with 1μM tetracycline and supplemented or not for 50 μM biotin were analyzed by immunoblot with antistreptavidin and anti-Flag antibodies. Anti-a-tubulin was used as a loading control. (C-D and F) WCE extracts of U2OS 2-6-5 cells transfected with the indicated GFP-(C and F) or mCherry-LacRnls- (D) proteins were analyzed by immunoblot with anti-GFP or anti-mCherry antibodies. GFP. Anti-a-tubulin was used as a loading control. (E) Representative images of U2OS 2-6-5 cells transfected with mCherry-LacRnls-FIGNL1 and the indicated GFP-C1orf112 constructs. GFP is used as control (Ctrl). Cells were fixed and mounted for confocal analyses. Scale bar = 5 μm (G) RT-qPCR for FIGNL1 was performed on U2OS cells. Expression was normalized against GADPH and reported as a percent relative to siCtrl. Data are represented as the mean ± SD (n = 3 independent experiments). Data were analyzed with an unpaired t-test with Welsh’s correction. (H) Viability assay in cells treated with DNA damage-inducing agent. U2OS cells were treated with the indicated siRNA. Forty-eight hours later, cells were either exposed to 2 μM cisplatin (Cis.), 150 μM formaldehyde (FA), 150 nM mitomycin C (MMC), 3 μM Talazoparib (BMN-673) for 48 hrs, to 2 hrs of 4 mM hydroxyurea (HU) followed by 46 hrs recovery, to 24 hrs of 1 mM HU followed by 24 hrs recovery, to 10 J/m^2^ UV or 10 Gy followed by 48 hrs recovery. Cells were counted with a hemacytometer and normalized against the number of cells quantified in the untreated condition. Data are represented as the mean ± SD (n = 4 independent experiments). Data were analyzed using a one-way Welch’s ANOVA test and Dunnett’s multiple comparison tests. **p<0.01, ***p<0.001, ****p<0.0001

**Figure S5.**
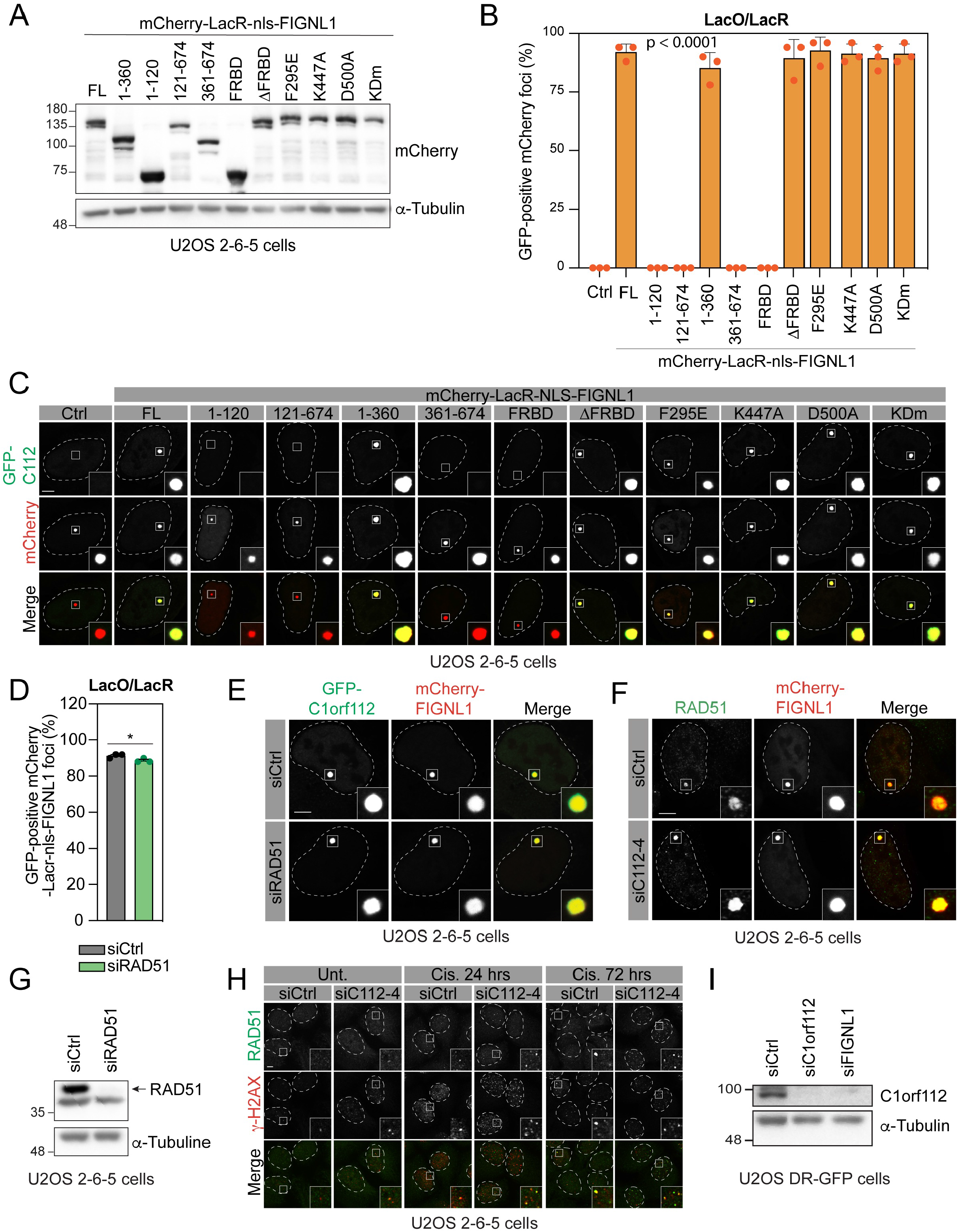
Expression levels and cellular localization of mCherry-LacRnls-FIGNL1 and GFP-C1orf112 constructs used in Fig. 5. (A) WCE of U2OS 2-6-5 cells transfected with the indicated mCherry-LacRnls-FIGNL1 constructs were analyzed by immunoblot with an anti-mCherry antibody. Anti-a-tubulin antibody was used as a loading control. KDm: K447A / D500A double mutant. (B) Quantification of mCherry-LacRnls constructs colocalizing with GFP-C1orf112 in U2OS 26-5 cells. Cells were treated and processed as described in Figure 4D. Data are represented as the mean ± SD (n = 3 independent experiments, 50 cells quantified per experiment in each condition). Data were analyzed with an unpaired t-test with Welsh’s correction. (C) Representative images of data from (B). U2OS 2-6-5 cells were transfected with indicated mCherry-LacR construct and then fixed and mounted for confocal analyses. Scale bar = 5 μm. (D) Quantification of the colocalization between mCherry-LacRnls-FIGNL1 and GFP-C1orf112 in U2OS 2-6-5 cells depleted or not for RAD51. Data are represented as the mean ± SD (n = 3 independent experiments, 50 foci quantified per experiment in each condition). Data were analyzed with an unpaired t-test. (E) Representative images of data quantified in (D). Cells were fixed and mounted for confocal analyses. Scale bar = 5 μm. (F) Representative images of the colocalization of mCherry-LacRnls-FIGNL1 with RAD51 in U2OS 2-6-5 cells depleted or not for C1orf112. Cells were fixed and processed for RAD51 immunofluorescence. Scale bar = 5 μm. (G) WCE of U2OS 2-6-5 cells depleted or not for RAD51 were analyzed by immunoblot with an anti-RAD51 antibody. Anti-a-Tubulin was used as a loading control. (H) Representative images of the data quantified in figure 5 E. Cells were fixed and processed for RAD51 (green) and g-H2AX (red) immunofluorescence. Scale bar = 5 μm. (I) WCE of U2OS DR-GFP cells depleted or not for C1orf112 or FIGNL1 were analyzed by immunoblot with an anti-C1orf112 antibody. Anti-a-Tubulin was used as a loading control. *p<0.05

**Figure S6.**
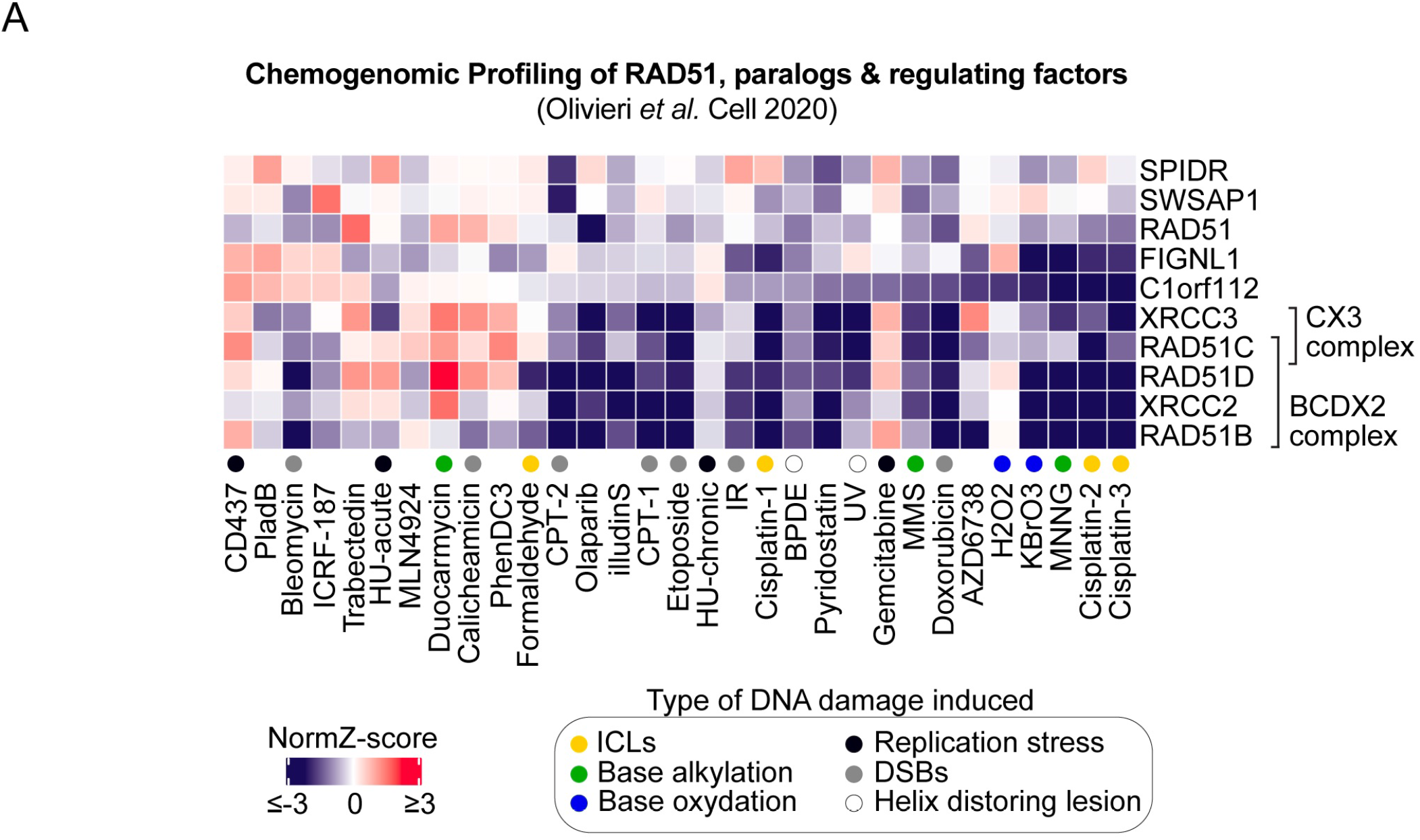
Chemogenomic profiling of C1orf112, FIGNL1 and literature-curated relevant genes. (A) Chemogenomic profiling using the NormZ-score from the CRISPR screens performed in RPE1 hTERT Cas9 p53^-/-^ cells with the indicated DNA damaging agents as detailed in^28^.

## Notes

### Competing Interest Statement

The authors have declared no competing interest.

