## Supplementary Tables S5-7 for "C1orf112 is a novel regulator of interstrand crosslink that decreases FIGNL1-RAD51 interaction"

**Table S5.** Primers, sgRNAs, shRNAs and siRNAs sequences used in this study.

| Name | Sequence 5'-3' | Used for[a] |
|---|---|---|
| sgCtrl (LacZ) | CCCGAATCTCTATCGTGCGG | KO in RPE1-hTERT Cas9 $p53^{-/-}$ |
| sgC1orf112-1 Exon 13 | TCCCTCACTGTTTGCTGAAC | KO in RPE1-hTERT Cas9 $p53^{-/-}$ |
| sgC1orf112-2 Exon 5 | TTATATGAAGGACTGAGGAG | KO in RPE1-hTERT Cas9 $p53^{-/-}$ |
| sgFANCA | CCACAGCATGCATGTCGGGA | KO in RPE1-hTERT Cas9 $p53^{-/-}$ |
| sgMRE11 | GCAATCATGACGATCCCACA | KO in RPE1-hTERT Cas9 $p53^{-/-}$ |
| sgFIGNL1-1 | GAAGACCCTGATGCACGCTG | KO in RPE1-hTERT Cas9 $p53^{-/-}$ |
| sgFIGNL1-2 | TTCTAAATGGGTAGGTGAGG | KO in RPE1-hTERT Cas9 $p53^{-/-}$ |
| shCtrl (scramble) (Addgene #1864) | CCTAAGGTTAAGTCGCCCTCG CTCGAGCGAGGGCGACTTAAC CTTAGG | KD in RPE1-hTERT WT and $p53^{-/-}$, and in IMR90 |
| shC1orf112-1 (TRCN0000163284) | GCTTCCTGACTATGTTCGTTT | KD in RPE1-hTERT WT and $p53^{-/-}$, and in IMR90 |
| shC1orf112-2 (TRCN0000165342) | GCAAGTTTCCTCCAAGCCTTT | KD in RPE1-hTERT WT and $p53^{-/-}$, and in IMR90 |
| siC1orf112-1 (D-020930-01) | CAUAGUGGCUCAUCUGAUA | KD in U2OS, HeLa DR-GFP, U2OS DR-GFP, U2OS SA-GFP & RPE1-hTERT $p53^{-/-}$ |
| siC1orf112-2 (D-020930-02) | AGACCUCGCUACUUAAAUU | KD in U2OS, U2OS DR-GFP, U2OS SA-GFP, HeLa DR-GFP & RPE1-hTERT $p53^{-/-}$ |
| siC1orf112-3 (D-020930-03) | GAAACGACAACCAGGAUAU | KD in U2OS, U2OS DR-GFP, U2OS SA-GFP, HeLa DR-GFP & RPE1-hTERT $p53^{-/-}$ |
| siC1orf112-4 (D-020930-04) | AGAGAUAGUUCCACAGUGU | KD in U2OS, U2OS 2-6-5, U2OS |

| | | |
|---|---|---|
| | | DR-GFP, U2OS SA-GFP, HeLa DR-GFP & RPE1-hTERT p53$^{-/-}$ |
| siFIGNL1-1 (D-019091-03) | GAGCAUGAAUCUUCUAGAA | KD in HeLa DR-GFP |
| siFIGNL1-2 (D-019091-17) | GCACAGAUAUUACGCAUUC | KD in U2OS, U2OS DR-GFP & HeLa DR-GFP |
| siCtIP (M-011376-00) | Smart pool<br>D-011376-01: GAGCAGACCUUUCUCAGUA<br>D-011376-02: GAAGUGAACAAGAUCAUUA<br>D-011376-03: CAACCAAGAUGUAUCCUUU<br>D-011376-04: GAAUAGGACUGAGUACGGU | KD in U2OS DR-GFP, HeLa DR-GFP & U2OS SA-GFP |
| siRAD51 (M-003530-04) | Smart pool:<br>D-003530-02: GAAGCUAUGUUCGCCAUUA<br>D-003530-05: GCAGUGAUGUCCUGGAUAA<br>D-003530-07: CCAACGAUGUGAAGAAAUU<br>D-003530-08: AAGCUAUGUUCGCCAUUAA | KD in U2OS, U2OS 2-6-5, U2OS DR-GFP & HeLa DR-GFP |
| siFANCA (custom) | GGACAGAUCUGCACGGCUC | KD in U2OS |
| siCtrl-1 (custom) | UGGUUUACAUGUCGACUAA | KD in U2OS and U2OS 2-6-5 |
| siCtrl-2 (D-001210-03) | UGGUUUACAUGUUUUCUGA | KD in U2OS and U2OS 2-6-5 |
| GAPDH FW | CAACGTGTCAGTGGTGGACC | RT, qPCR |
| GAPDH RV | TCGTTGAGGGCAATGCCAGC | RT, qPCR |
| FIGNL1 FW | ACCAGCCGCAGGTGAAAAC | qPCR |
| FIGNL1 RV | ACAATGCTCCTTGATGCTGC | qPCR |
| TKO library outer-PCR primer FW | AGGGCCTATTTCCCATGATTCCTT | PCR |
| TKO library outer-PCR primer RV | TCAAAAAAGCACCGACTCGG | PCR |
| TKO library inner-PCR primer trueseq i5 1 | AATGATACGGCGACCACCGAGATC<br>TACACTATAGCCTACACTCTTTCCC<br>TACACGACGCTCTTCCGATCTTGTG<br>GAAAGGACGAGGTACCG | PCR |
| TKO library inner-PCR primer trueseq i5 2 | AATGATACGGCGACCACCGAGATC<br>TACACATAGAGGCACACTCTTTCC<br>CTACACGACGCTCTTCCGATCTTGT<br>GGAAAGGACGAGGTACCG | PCR |
| TKO library inner-PCR primer trueseq i5 3 | AATGATACGGCGACCACCGAGATCT<br>ACACCCTATCCTACACTCTTTCCCTA<br>CACGACGCTCTTCCGATCTTGTGGAA<br>AGGACGAGGTACCG | PCR |

| | | |
|---|---|---|
| TKO library inner-PCR primer trueseq i5 4 | AATGATACGGCGACCACCGAGATCTACACGGCTCTGAACACTCTTTCCCTACACGACGCTCTTCCGATCTTGTGGAAAGGACGAGGTACCG | PCR |
| TKO library inner-PCR primer trueseq i7 1 | CAAGCAGAAGACGGCATACGAGATCGAGTAATGTGACTGGAGTTCAGACGTGTGCTCTTCCGATCTATTTTAACTTGCTATTTCTAGCTCTAAAAC | PCR |
| TKO library inner-PCR primer trueseq i7 2 | CAAGCAGAAGACGGCATACGAGATTCTCCGGAGTGACTGGAGTTCAGACGTGTGCTCTTCCGATCTATTTTAACTTGCTATTTCTAGCTCTAAAAC | PCR |
| TKO library inner-PCR primer trueseq i7 3 | CAAGCAGAAGACGGCATACGAGATAATGAGCGGTGACTGGAGTTCAGACGTGTGCTCTTCCGATCTATTTTAACTTGCTATTTCTAGCTCTAAAAC | PCR |
| TKO library inner-PCR primer trueseq i7 4 | CAAGCAGAAGACGGCATACGAGATGGAATCTCGTGACTGGAGTTCAGACGTGTGCTCTTCCGATCTATTTTAACTTGCTATTTCTAGCTCTAAAAC | PCR |
| TKO library inner-PCR primer trueseq i7 5 | CAAGCAGAAGACGGCATACGAGATTTCTGAATGTGACTGGAGTTCAGACGTGTGCTCTTCCGATCTATTTTAACTTGCTATTTCTAGCTCTAAAAC | PCR |
| TKO library inner-PCR primer trueseq i7 6 | CAAGCAGAAGACGGCATACGAGATACGAATTCGTGACTGGAGTTCAGACGTGTGCTCTTCCGATCTATTTTAACTTGCTATTTCTAGCTCTAAAAC | PCR |
| C1orf112 FL attb1 cloning FW | GGGGACAAGTTTGTACAAAAAAGCAGGCTTCATGTTTTTACCTCATATGAACCACC | PCR |
| C1orf112 FL attb1 cloning RV | GGGGACCACTTTGTACAAGAAAGCTGGGTTTCACCCTAGAGTATGTATGTAACGT | PCR |
| C1orf112 NTD quickchange oligo FW | GGGGACAAGTTTGTACAAAAAAGCAGGCTTCATG CATGCATTTCATGCCAATACTTGGA | PCR |

| | | |
|---|---|---|
| C1orf112 NTD quickchange oligo RV | GGGGACCACTTTGTACAAGAAAGCTGG GTTTCACCCTAGAGTATGTATGTAACGT | PCR |
| C1orf112 ΔCTD quickchange oligo FW | GTTCGCTGAGGGAACAAATCATGAAGA GATATAGCCATAGTGTCTCAGTTCTGA | PCR |
| C1orf112 ΔCTD quickchange oligo RV | TCAGAACTGAGACACTATGGCTATATC TCTTCATGATTTGTTCCCTCAGCGAAC | PCR |
| C1orf112 ΔWCF quickchange oligo FW | TTTGTTAGCTATGGATGCACTTGCTCGA TATGGGACTG | PCR |
| C1orf112 ΔWCF quickchange oligo RV | CAGTCCCATATCGAGCAAGTGCATCCAT AGCTAACAAA | PCR |
| FIGNL1 1-120 & 1-360 oligo FW | GGGGACAAGTTTGTACAAAAAAGCAGG CTCCATGCAGACCTCCAGCTCTA | PCR |
| FIGNL1 1-120 oligo RV | GGGGACCACTTTGTACAAGAAAGCTGGG TCCTATTGCATCATCTTCTGTACACTACTC | PCR |
| FIGNL1 1-360 oligo RV | GGGGACCACTTTGTACAAGAAAGCTGGG TCCTAAGGCTTACATTGCATTCCT | PCR |
| FIGNL1 361-674 oligo FW | GGGGACCACTTTGTACAAGAAAGCTGGG TCCTATTGCATCATCTTCTGTACACTACTC | PCR |
| FIGNL1 121-674 oligo FW | GGGGACAAGTTTGTACAAAAAAGCAGGC TCCATGGCTGGCAAAAAATTCAAAGA | PCR |
| FIGNL1 361-674 & 121-674 oligo RV | GGGGACCACTTTGTACAAGAAAGCTGGG TCTTACTTTCCACAACCAAAAGTTTT | PCR |
| FIGNL1 FRBD (aa) oligo FW | GGGGACAAGTTTGTACAAAAAAGCAGGC TCCATGTTTAAAACTGCAAAAGAACAATT AT | PCR |
| FIGNL1 FRBD oligo RV | GGGGACCACTTTGTACAAGAAAGCTGGG TCCTATATAGGAGGAACAAACTTTCC | PCR |
| FIGNL1 ΔFRBD oligo FW | CCCAAGCAAGATGGGGGAGA | PCR |
| FIGNL1 ΔFRBD oligo RV | TGTAGGCAGGCTGCTATCCTCC | PCR |
| FIGNL1 F295E oligo FW | CCTACAGAGAAAACTGCAAAAGAAC | PCR |
| FIGNL1 F295E oligo RV | CAGGCTGCTATCCTCCTTTG | PCR |
| FIGNL1 K447A oligo FW | GGTGCAACTCTAATTGGCAAG | PCR |
| FIGNL1 K447A oligo RV | AGTCCCAGGAGGACCAAAG | PCR |
| FIGNL1 D500A oligo FW | ATTGCCGAAATTGATTCCTTG | PCR |

| | | |
|---|---|---|
| FIGNL1 D500A oligo RV | AAATATCACAGCTGGTTGCTG | PCR |

[a] KD: knockdown

**Table S6.** Antibodies used in this study

| **Primary antibodies** | | |
|---|---|---|
| **Target** | **Catalog number** | **Used for** |
| Rabbit anti-53BP1 | NB100-304 (Novus) | IF |
| Rabbit anti-BRCA1 | 07-434 (Millipore) | IF |
| Mouse anti-BRCA2 | OP95 (Calbiochem, Millipore) | IF |
| Mouse anti-Cas9 | 14697 (Cell signaling) | |
| Human anti-centromere (CREST) | HCT-0100 (Immunovision) | IF |
| Rabbit anti-mCherry | NBP2-25157 (Novus) | WB |
| Mouse anti-Cyclin A1 | 611269 (BD Bioscience) | IF |
| Rabbit anti-C1orf112 | ab121774 (Abcam) | WB |
| Rabbit anti-FANCA | A301-980A-M (Bethyl) | WB |
| Rabbit anti-FANCD2 | NB100-182 (Novus) | IF |
| Mouse anti-FLAG | M2 (Sigma) | WB |
| Mouse anti-GFP IgG1κ (clones 7.1 and 13.1) | 11814460001 (Sigma) | WB |
| Mouse anti-phospho-Histone H2A.X (Ser139) (γ-H2AX) | 05-636 (Millipore) | IF |
| Rabbit anti-Lamin-B | Ab16048 (Abcam | IF |
| Rabbit anti-RAD51 | 70-001 (Bio Academia) | IF |
| Anti-Streptavidin HRP conjugated | RPN1231V (GE Healthcare) | WB |
| Mouse anti-α-Tubulin | CP06 (Sigma) | WB |
| **Secondary antibodies** | | |
| **Target** | **Catalog number** | **Used for** |
| Alexa Fluor 488 goat anti-rabbit | A-11034 (Thermo Fisher Scientific) | IF |
| Alexa Fluor 555 goat anti-rabbit | A-21428 (Thermo Fisher Scientific) | IF |
| Alexa Fluor 647 goat anti-rabbit | A-21244 (Thermo Fisher Scientific) | IF |
| Alexa Fluor 488 goat anti-mouse | A-11029 (Thermo Fisher Scientific) | IF |
| Alexa Fluor 555 goat anti-mouse | A-21424 (Thermo Fisher Scientific) | IF |
| Alexa Fluor 647 goat anti-mouse | A-21236 (Thermo Fisher Scientific) | IF |
| Alexa Fluor 647 donkey anti-human | A-21445 (Thermo Fisher Scientific) | IF |
| Goat anti-rabbit HRP | 7074 (Cell Signaling) | WB |
| Sheep anti-mouse-HRP | A-9044 | WB |

**Table S7**. Plasmids used in this study

| Plasmid | Source |
|---|---|
| lentiCas9-Blast | Addgene #52962 |
| Toronto KnockOut (TKO) CRISPR library – v1 | Addgene #1000000069 |
| pKLV2-U6gRNA5(BbsI)-PGKpuro2AmCherry | Addgene #67977 |
| pKLV2-U6gRNA5(BbsI)-PGKpuro2ABFP | Addgene #67991 |
| pOG44 | Invitrogen™ V600520 |
| pDEST-pcDNA5-FLAG-miniturbo | This study |
| FLAG-miniturbo-C1orf112 | This study |
| pDEST-pcDNA5-FRT-TO-mCherry-LacRnls | Described in (Orthwein *et al*, 2015) |
| mCherry-LacRnls C1orf112 FL (1-854) | This study |
| mCherry-LacRnls FIGNL1 1-674 | This study |
| mCherry-LacRnls FIGNL1 1-120 | This study |
| mCherry-LacRnls FIGNL1 121-674 | This study |
| mCherry-LacRnls FIGNL1 1-360 | This study |
| mCherry-LacRnls FIGNL1 361-674 | This study |
| mCherry-LacRnls FIGNL1 FRBD (295-344) | This study |
| mCherry-LacRnls FIGNL1 ΔFRBD (Δ295-344) | This study |
| mCherry-LacRnls FIGNL1 F295E | This study |
| mCherry-LacRnls FIGNL1 K447A | This study |
| mCherry-LacRnls FIGNL1 D500A | This study |
| mCherry-LacRnls FIGNL1 KDm (K447A/D500A) | This study |
| pDEST-pcDNA5-FRT-TO-eGFP | Described in (Escribano-Díaz *et al*, 2013) |
| GFP-C1orf112 FL (1-854) | This study |
| GFP-C1orf112 NTD (1-175) | This study |
| GFP-C1orf112 ΔCTD (1-735) | This study |
| GFP-C1orf112 ΔWCF (461-463) | This study |
| GFP-FIGNL1 FL (1-674) | This study |
| pCBASceI | Addgene #26477 |

| Tool name | Reference | Source |
|---|---|---|
| MAGecK 0.5.9.5 | https://sourceforge.net/p/mageck/wiki/Home/ | (Li *et al*, 2014) |
| DrugZ | https://github.com/hart-lab/drugz | (Colic *et al*, 2019) |
| R-Studio 22.07.1 | https://www.r-project.org/ | R software |
| GSEA 4.3.0 | https://www.gsea-msigdb.org/gsea/index.jsp | (Subramanian *et al*, 2005; Mootha *et al*, 2003) |

| | | |
|---|---|---|
| Graphpad-Prism 9.3.1 | https://www.graphpad.com/scientific-software/prism/ | GraphPad |
| Snapgene 6.0.6 | https://www.snapgene.com | Dotmatics |
| FlowJo 10.7.2 | https://www.flowjo.com | FlowJo LLC |
| Fiji – ImageJ 2.3.0 | https://imagej.net/Fiji | (Schneider *et al*, 2012) |
| Adobe Photoshop 22.1.0 | https://www.adobe.com/products/illustrator.html | Adobe |
| Adobe Illustrator 25.0.1 | https://www.adobe.com/products/photoshop.html | Adobe |
